# Efficient Generation of Expandable Dorsal Forebrain Neural Rosette Stem Cell Lines

**DOI:** 10.1101/2025.05.27.656305

**Authors:** Signe Emilie Dannulat Frazier, Kristian Honnens de Lichtenberg, Elham Jaberi, Charlotte Bertelsen, Simone Møller Jensen, Andreas Wrona, Nicolaj Strøyer Christophersen, Mie Kristensen, J. Carlos Villaescusa

**Affiliations:** Novo Nordisk, Cell Therapy R&D, DK-2760 Måløv, Denmark; Department of Pharmacy, University of Copenhagen, Copenhagen 2100, Denmark

## Abstract

Neural stem cells (NSCs) represent an interesting option for developing *in vitro* disease models and drug screening assays due to their differentiation capacity into neurons and glial cells. Additionally, NSCs are under investigation in on-going clinical trials for treatment of various human neurological disorders. NSCs can be isolated from the central nervous system or derived *in vitro* from human pluripotent stem cells (hPSCs). However, the current methods for generating NSCs typically include a phase of neural rosette formation and subsequent manual isolation of these tiny structures. As this is a laborious process characterized by operator-dependent variability and scalability challenges, there is a pressing need to develop optimized and scalable protocols to obtain pure NSC populations. In this study, we present a new method for generating highly pure and expandable dorsal forebrain FOXG1^+^OTX2^+^TLE4^+^SOX5^+^ neural rosette stem cell (NRSC) lines without the necessity for manual isolation of rosette structures. Our findings demonstrate the reproducibility of this protocol through the characterization of different NRSC lines over multiple passages, highlighting the robustness of the process. These NRSCs can be expanded for at least 12 passages without compromising their rosette-formation capacity or their initial dorsal forebrain identity. Furthermore, we show the differentiation capacity of these NRSCs to generate pure populations of TUBB3^+^ neurons, and under specific conditions, their ability to differentiate into early glial progenitor cells including GFAP^+^ astrocytes and O4^+^ oligodendrocytes. Collectively, these results show the capabilities of our protocol to generate an expandable NRSC population suitable for *in vitro* disease modeling and drug screening, while also suggesting a viable strategy for scalable NRSC production for clinical application.

## Introduction

NSCs are recognized as important tools for the development of drug screening platforms and *in vitro* disease models, particularly in the context of neurological disorders. Their unique capacity for self-renewal and differentiation into different types of neurons, astrocytes, and oligodendrocytes positions them as potential candidates for clinical translation. NSCs contribute to neuroprotection through mechanisms that include cellular replacement and the secretion of neuroprotective factors (Alonso-Olivares *et al*., 2024; Hudáčová, 2021).

The generation of NSCs can be accomplished through direct isolation from the central nervous system or via differentiation of hPSCs. Currently, the most common methods utilize the formation of neural rosettes followed by manual isolation to increase purity, which introduces significant variability and challenges in scalability. As highlighted by several studies, this manual approach can be labor-intensive and inconsistent, reflecting the need for standardized protocols that enhance reproducibility across multiple passages (Dady *et al*., 2022; Verrier *et al*., 2018; Hříbková *et al*., 2018). The difficulties encountered in the isolation of neural rosettes emphasize the necessity of refining methodologies to ensure consistency and reliability in NSC production.

Recent explorations have focused on alternative protocols that avoid the manual selection step of neural rosettes. However, these newer approaches often lead to heterogeneous NSC populations that struggle with stability during passaging (D’Aiuto *et al*., 2022; Knight *et al*., 2018). The optimization of protocols to facilitate scalable production of NSCs is therefore important to ensure their clinical applicability.

This study employs dynamic suspension culture coupled with single SMAD inhibition to generate neural rosette stem cells (NRSCs) with high purity and stability over extended passages. Furthermore, we have investigated the multipotency of these cells through differentiating them into functional neurons, astrocytes, and oligodendrocytes, thus ensuring their differentiation potential.

Our research aims to overcome the limitations associated with traditional protocols by introducing an innovative strategy for generating highly pure dorsal forebrain populations of NRSCs characterized by markers such as FOXG1, OTX2, TLE4, and SOX5 without the necessity for manual isolation. Notably, these NRSCs maintain their capacity for rosette formation across more than 12 passages, simultaneously retaining their inherent identity associated with the dorsal forebrain (Kim *et al*., 2024; Birtele *et al*., 2023). This protocol not only enhances the reproducibility of NSC generation but also aligns with the growing necessity for efficient and standardized procedures in stem cell research, paving the way for new *in vitro* modelling and future innovative therapeutic strategies.

## Results

### Generation of high-purity populations of NRSCs without the need of manual selection

Most protocols for generating NSCs from hPSCs involve a manual selection step, where neural rosettes are either manually picked or selected using agents to achieve purer populations (Zhang *et al*., 2001; Elkabetz *et al*., 2008; Koch *et al*., 2009; Bohaciakova *et al*., 2019). In contrast, protocols that do not perform manual selection of rosettes often demonstrate high heterogeneity or variability in marker composition (Ebert *et al*., 2013; Wen and Jin, 2014; Fedorova *et al*., 2019).

This study presents a protocol for deriving highly pure populations of NRSCs that are expandable and identified by the continuous formation of neural rosettes after multiple passages. Notably, our process does not require the manual selection step of neural rosettes to efficiently convert human embryonic stem cells (hESCs) into NRSCs.

The protocol is divided into three different phases: neuroectoderm induction, rosette formation, and NRSC line establishment (**Fig. 1A**). hESCs were dissociated into a suspension of single cells and placed into non-adhesive plates, where they were kept in static culture for 24 hours to allow for the natural formation of floating cell spheres. It is important not to disturb the cultures during this phase to avoid aggregation of the cell spheres. After 24 hours, the culture conditions were switched from static to dynamic culture by transferring the spheres onto an orbital shaker set to 40 RPM with a 3 second vibration step every 15 seconds. The cells were exposed to single SMAD inhibition from day 0 to 10 to induce neuroectoderm formation. We evaluated various SMAD signaling inhibitors and found that the small molecule RepSox was the most effective in promoting the formation of rosette structures and enhancing cell purity for NES (Nestin) and OTX2 (data not shown).

**Figure 1.**
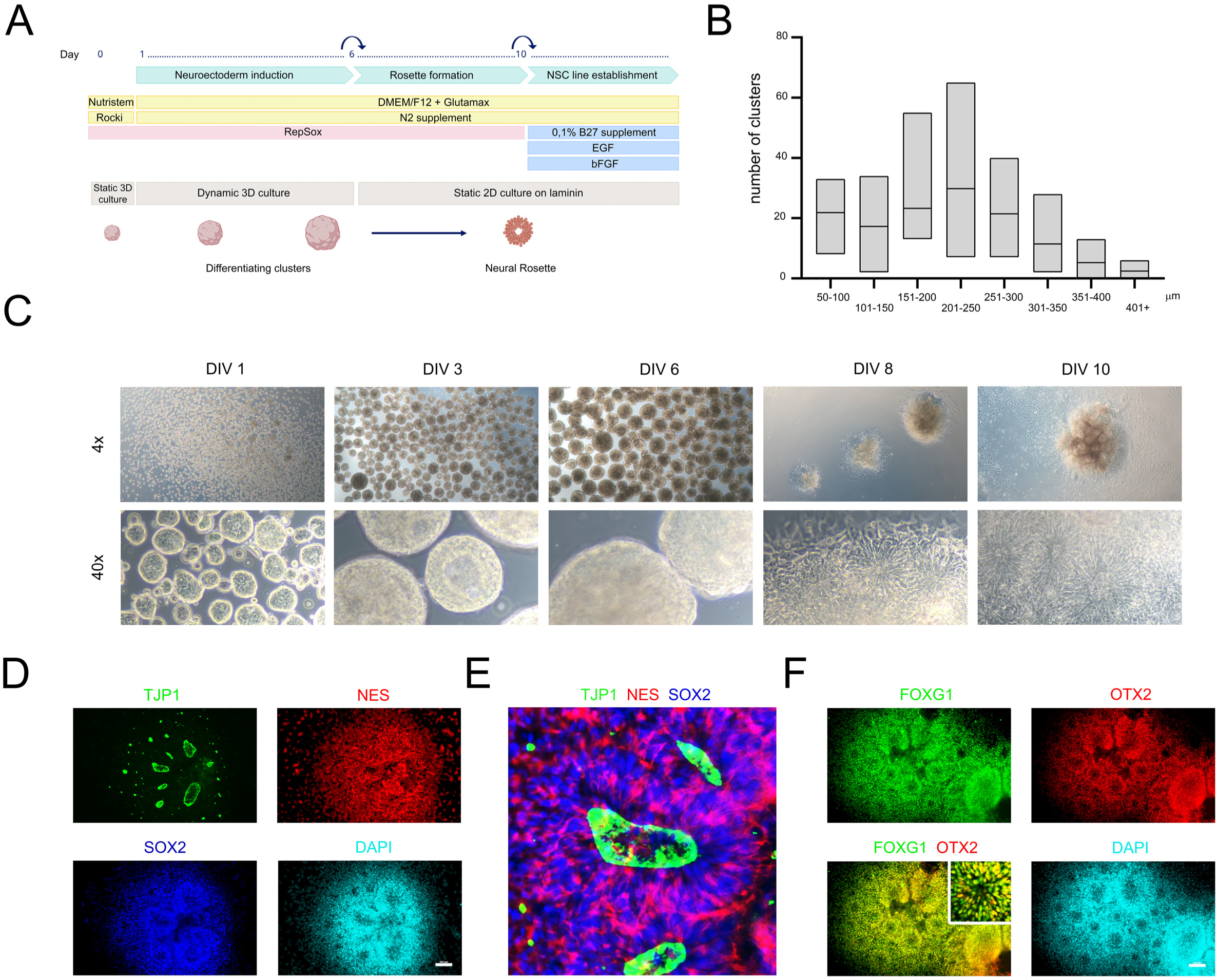
Generation of high purity NRSCs with no manual selection of rosettes. **(A)** Schematic representation of protocol for the generation of highly pure populations of NRSCs with no manual selection. **(B)** Size distribution of differentiating clusters measured on DIV 6 before seeding in 2D. Box represents minimum and maximum values, line represents mean, N=5. **(C)** Representative brightfield images captured on DIV (days *in vitro*) 0, 1, 3, 6, 8 and 10 in 4x magnification and 40x magnification. **(D)** Representative immunofluorescence images of DIV10 NRSCs positive for the common NSC markers NES and SOX2. **(E)** Higher magnification of a neural rosette. **(F)** NRSCs were also positive for the forebrain markers FOXG1 and OTX2. The formation of neural rosettes is visualized by the redistribution of TJP1 into the lumen of the rosettes. Scalebar = 100 µm.

By day 6, most clusters ranged from 100 to 400 µm in size (**Fig. 1B**) and exhibited a morphologically smooth surface, with the onset of cavitation visible via brightfield imaging (**Fig 1C**). Clusters were then seeded in 2D on laminin-coated plates for an additional 4 days, during which they initiated flattening into a cell monolayer of neural rosettes, observable by brightfield imaging (**Fig. 1C**). On day 10, the cells were positive for the typical NSC marker NES and the proliferation marker SOX2 (**Fig. 1D**). Furthermore, we observed re-distribution of the tight junction protein TJP1 (also known as zonula occludens-1, ZO1) into the lumen of the rosettes, indicating a polarized organization (**Fig. 1E**).

The NRSCs generated with this protocol exhibit a defined forebrain regional identity, characterized by the rostral markers FOXG1 and OTX2 (**Fig. 1F**). On day 10, the flattened clusters were enzymatically dissociated into single cells and seeded at a density of 1.5 million cells/cm² for expansion. It is crucial to maintain a high density at these early passages to enhance cell survival and promote the reformation of small neural rosette structures. The NRSCs were cultured for 3 days before being passaged again at a density of 1.5 million cells/cm^2^. This process was repeated 3 times to obtain a homogeneous population of rosette-forming NRSCs, and to establish the NRSC line, which can subsequently be passaged at lower seeding densities of down to 0.5 million cells/cm^2^ for up to 12 passages, while preserving their defining characteristics and regional identity. NRSCs could be dissociated into single cells and cryopreserved, with an average viability above 80% upon thawing (**Suppl. Fig. 1A**). This simplified approach demonstrates the generation of highly pure and expandable NRSCs within just 10 days.

### Reproducibility of NRSC line generation and maintenance of homogeneity across passaging

The ability to generate stable and comparable NSC lines through multiple passages is essential for the feasibility of large-scale manufacturing. We evaluated the reproducibility of our differentiation protocol by establishing two distinct neural rosette-forming NSC lines from the same parental hPSC line. Both NRSC lines exhibited morphologically similar characteristics, primarily comprising small, proliferating rosette-forming NSCs that could be expanded for at least 12 passages (**Fig. 2A and Suppl. Fig. 2A**).

**Figure 2.**
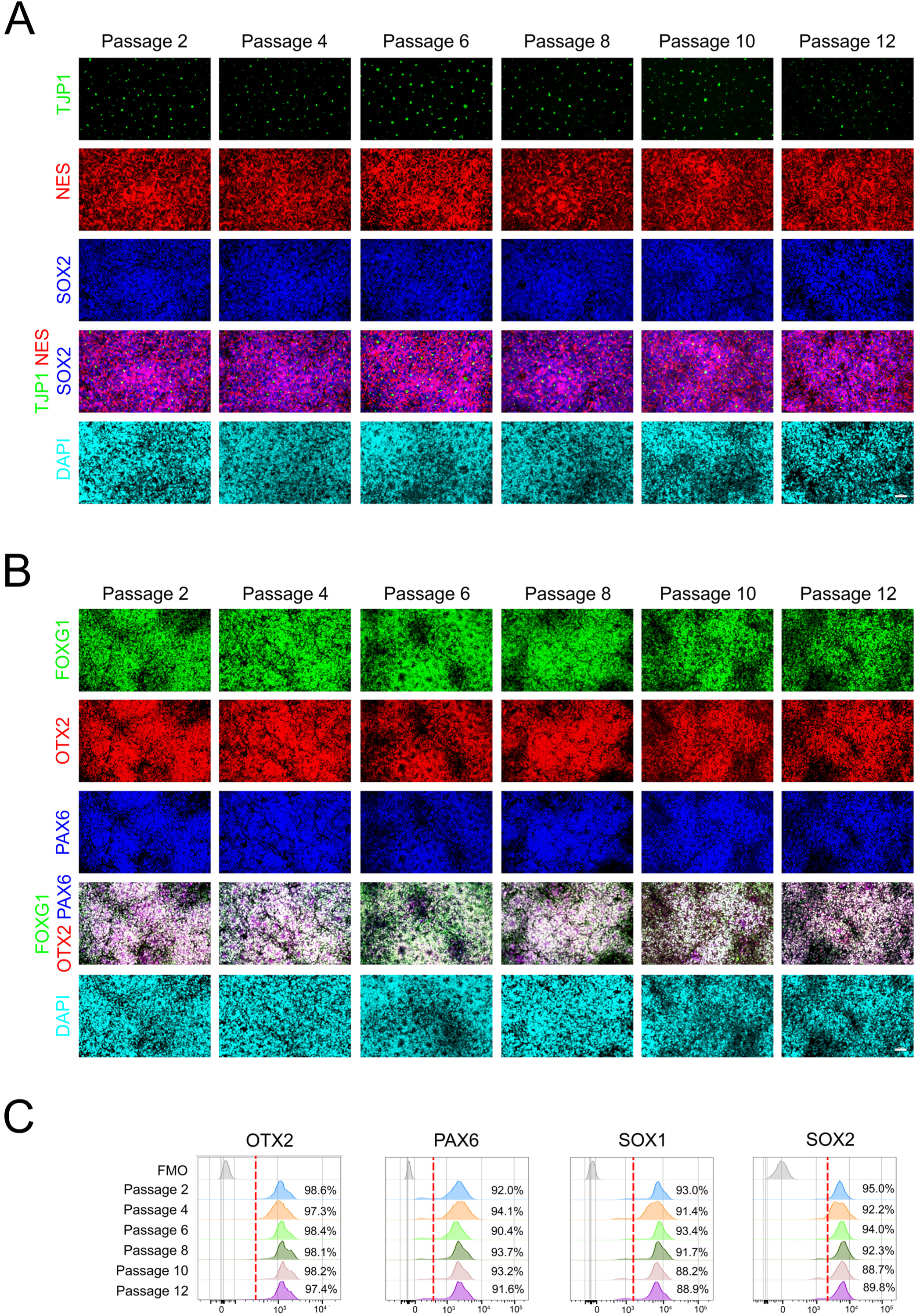
NRSCs maintain stable marker expression through extended passaging. **(A)** Representative immunofluorescence images showing cells positive for NSC markers NES and SOX2 at passages 2, 4, 6, 8, 10 and 12. TJP1 is used to highlight the lumen of neural rosettes. All images were taken in 20X magnification, scalebar = 50 µm. The different passages were imaged at different timepoints. **(B)** Representative immunofluorescence images showed NRSC positive for regional forebrain markers FOXG1 and OTX2 and NSC marker PAX6. All images were taken in 20X magnification, scalebar = 50 µm. The different passages were imaged at different timepoints. **(C)** Flow cytometric analysis of NSCs at passages 2, 4, 6, 8, 10 and 12. The cells showed stable marker expression through passaging of NSC markers SOX2, PAX6, neuroectoderm marker SOX1 and forebrain marker OTX2. Red dashed lines delineate the gating regions corresponding to the displayed percentages. FOM, Fluorescence Minus One.

The neural rosettes displayed small, centralized lumens, confirmed by TJP1 positivity of the neural rosette lumen, as well as a radial organization, with NSCs expressing the markers NES and SOX2. Assessment of the TJP1+ rosette lumens demonstrated that the NRSC populations consistently reformed rosettes throughout passaging. This pattern was maintained at least until passage 12, when we concluded the NRSC expansions. The NRSCs proliferated at a rate of approximately every 3-4 days (data not shown).

Moreover, our NRSCs were positive for the anterior identity markers FOXG1 and OTX2, along with PAX6, collectively indicating their forebrain identity (**Fig. 2B and Suppl. Fig. 2B**). Across passages 2 to 12, no significant changes were detected in the proportion of positive cells for FOXG1, OTX2, or PAX6, as determined by immunofluorescence. This was further validated by flow cytometry, which revealed that OTX2+ cells constituted over 95% of the population from passages 2 to 12, while PAX6+ cells exceeded 90%. Additionally, the early neuroepithelial marker SOX1 was present in over 88% of cells, and SOX2 positivity remained above 89% at passage 12 (**Fig. 2C**).

To further assess reproducibility across different hPSC lines, we applied the same 10-day differentiation protocol (**Fig. 1A**) to another hESC line. The results demonstrated that rosette-forming NRSC cultures with consistent morphological and immunocytochemical characteristics could be derived from different hPSC lines (**Suppl. Fig.3**). These findings indicate that our protocol reliably and consistently generates stable and pure NRSC populations that retain their identity over extended passages.

### Transcriptomic analysis reveals stability and forebrain identity of NRSCs during passaging

We proceeded to analyze our NRSCs at the transcriptomic level. Single-cell RNA sequencing (scRNA-seq) was conducted at both passage 2 and passage 8 to investigate cellular identity and any potential changes occurring during passaging.

The expression levels of the common NSC markers *SOX2*, *NES*, and *DACH1*, remained relatively stable between passages 2 and 8 (**Fig. 3A**). However, *PAX6* expression decreased by approximately 25% from passage 2 to passage 8 (**Fig. 3A**), a finding that did not correlate with the flow cytometry and immunofluorescence data (**Fig. 2B and C**). A possible explanation for this discrepancy could be the negative autoregulation of *PAX6* expression, as reported previously (Manuel *et al*., 2007). The high levels of *DACH1*, recognized as a marker of neural rosette progenitors *in vitro* (Elkabetz *et al*., 2008), further confirmed the identity of our NRSCs.

**Figure 3.**
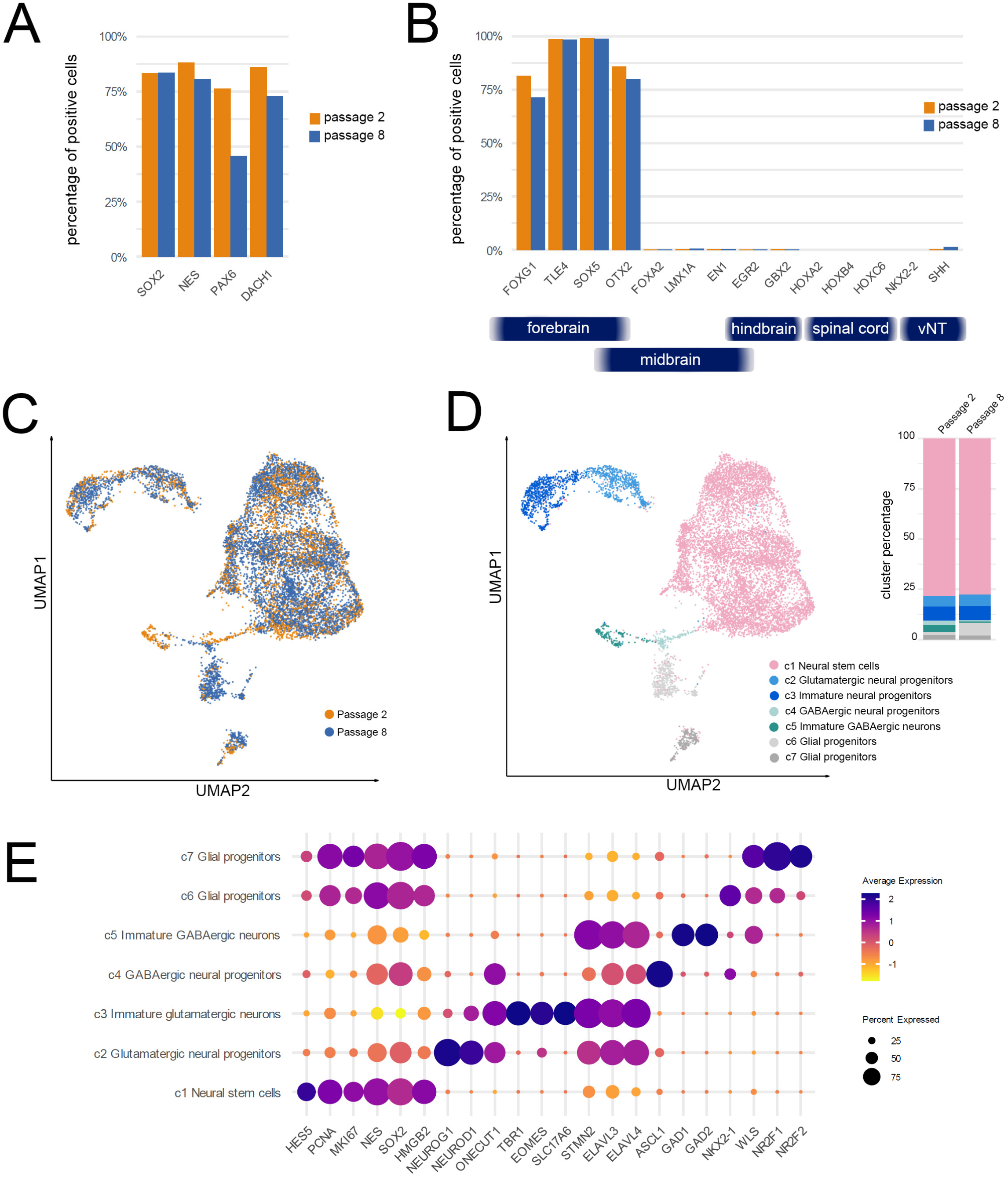
scRNAseq reveals the presence of mainly forebrain NRSCs in cultures. **(A)** Percentage of NRSCs expressing classic NSC stem cell markers at passage 2 and 8. Passage 2 shown in orange and passage 8 in blue. **(B)** Percentage of NRSCs expressing markers of regional identity in the developing brain. NRSCs show high and consistent expression of forebrain regional markers *FOXG1*, *TLE4*, *SOX5*, *OTX2*, with no midbrain, hindbrain, spinal cord or vNT (ventral neural tube) contaminants in both passage 2 and 8. Passage 2 shown in orange and passage 8 in blue. **(C)** UMAP of passage 2 and 8 NRSCs integrated, showing no major differences in the cell-types present. Passage 2 shown in yellow and passage 8 in blue. **(D)** UMAP of NRSC cultures, revealed large cluster of proliferating NRSC and minor populations of slightly more mature cells. Annotation is based on known markers highly expressed within each cluster. UMAP shows integrated data from passage 2 and 8. Percentage of cells in each cluster is shown as bar plot. **(E)** Dotplot visualizing the expression levels of NSC-, neural progenitor-, neuronal- and glial progenitor markers in the different clusters.

Next, we examined the regional identity of our NRSCs by assessing the expression of *FOXG1* and *OTX2*, which collectively confirm the forebrain identity of our NRSCs (**Fig. 3B**). The expression levels of these anterior markers appeared stable between passages 2 and 8, reinforcing a consistent forebrain identity. This stability is aligned with the immunofluorescence (**Fig. 2B**) and flow cytometry results (**Fig. 2C**). Additionally, we observed consistent expression of *TLE4* and *SOX5*, both known to be present in the dorsal forebrain during development (Azim *et al*., 2009; Galazo, Sweetser and Macklis, 2023). In contrast, midbrain markers such as *FOXA2*, *LMX1A*, and *EN1* were absent in our NRSC cultures at both passages. Moreover, specific markers for the embryonic hindbrain, such as *EGR2* and *GBX2*, as well as the spinal cord markers *HOXA2*, *HOXB4*, and *HOXC6*, were not detected in our NRSCs. Interestingly, *NKX2-2*, primarily expressed in the ventral regions of the developing forebrain, was completely absent from our NRSC cultures. Furthermore, *SHH*, a marker for the ventral neural tube, was also absent, supporting the idea that our NRSCs corresponded to a dorsal forebrain identity (**Fig. 3B**).

Overall, it is notable that our NRSCs exhibit a remarkable regional purity, consisting almost entirely of dorsal forebrain NRSCs, with no contamination from other brain regions. Consistent examination of the Uniform Manifold Approximation and Projection (UMAP) plots for passages 2 and 8 revealed similar patterns, and integrating the datasets showed no major changes in the existing cell clusters (**Fig. 3C**).

The clusters identified in the UMAP analyses were annotated based on the expression of known markers. A substantial proportion of NRSCs, cluster c1, expressed genes associated with common NSCs, including *NES*, *SOX2*, *PAX6*, *OTX2*, *HES5*, *MSI1*, *TTYH1*, *PRDM16*, *VIM*, *MKI67*, and *PCNA* (**Fig. 3D and E**, **and Suppl. Fig. 4A**). These NSC markers (Kaneko *et al*., 2000; Chuikov *et al*., 2010; Zhang and Jiao, 2015; Shimada *et al*., 2017; Kim *et al*., 2018) facilitate the maintenance and differentiation of NSCs, whereas *MKI67* and *PCNA* are expressed by proliferating cells in the brain and other tissues (Zhang and Jiao, 2015). This cluster represented 78% of the total cells at passage 2 and 77% at passage 8, demonstrating high purity and stability of our NRSC culture over multiple passages.

In addition to the NRSCs, we identified three major trajectories. Cluster c2 and c3 were associated with trajectories toward immature glutamatergic neuron differentiation (**Fig. 3D**). Cluster c2 expressed neural progenitor markers *NEUROD1* and *NEUROG1* (**Fig. 3E**, **Suppl. Fig. 4B**). *NEUROD1* has been implicated in the fate specification of developing glutamatergic neurons (Hevner *et al*., 2006), while *NEUROG1* promotes glutamatergic differentiation (Reyes *et al*., 2008). In contrast, cluster c3 expressed more mature glutamatergic neuronal markers, including *TBR1*, *EOMES*, and *SLC17A6* (**Fig. 3E and Suppl. Fig. 4B**). *EOMES* is recognized as a marker for intermediate progenitor cells committed to a glutamatergic fate, while *TBR1* is known to be expressed by all post-mitotic glutamatergic neurons (Englund *et al*., 2005; Hevner *et al*., 2006), and *SLC17A6* serves as a well-established marker for glutamatergic axon terminals (Nakamura *et al*., 2005).

Cluster c4 was linked to trajectories toward immature GABAergic neuron differentiation (**Fig. 3D**), expressing progenitor markers *ASCL1*, *NKX2.1*, and *DLX2* (**Fig. 3E**, **Suppl. Fig. 4C**). Both *ASCL1* and *DLX2* have been implicated in GABAergic neuron fate specification (Casarosa, Fode and Guillemot, 1999; Virolainen *et al*., 2012), while *NKX2.1* is a marker of the medial ganglionic eminence (MGE), giving rise to specific GABAergic neurons and glial cells (Nóbrega-Pereira *et al*., 2008). This suggests that cluster c4 may represent a common progenitor for both GABAergic and glial trajectories. Cluster c5 was characterized by the presence of well-known GABAergic markers, including *GAD1/2* and *DLX2*.

Finally, clusters c6 and c7 comprised a glial progenitor pool characterized by *GLIS3*, *NKX2.1*, *NR2F1/2*, and *WLS* (Fig. 3E and Suppl. Fig. 3D). Both c6 and c7 were enriched in *GLIS3*, a gene associated with the neuro-glial switch (Van Bruggen *et al*., 2021). Additionally, c6 expressed the oligodendrocyte lineage marker *NKX2.1*, while c7 exhibited high expression of *NR2F1/2* and *WLS*, which are crucial for glial commitment (Belgacem *et al*., 2016).

Together, these results indicate that our cell culture predominantly consists of a large pool of proliferating NRSCs with smaller populations differentiating into neural and glial lineages. Notably, it was possible to maintain these differentiating populations without significant changes over passages.

These findings demonstrate that our protocol reliably produces a highly pure population of proliferating dorsal forebrain NRSCs that maintain stable marker expression across several passages with minimal differentiation.

### Neurogenic potential of forebrain NRSCs to differentiate into functional neurons

We investigated the ability of our forebrain NRSCs to differentiate into neurons. Notably, our NRSCs expressed the transcription factor gene *GLI3* (**Suppl. Fig. 4A**), which is crucial for the dorsal-ventral patterning in cooperation with SHH signaling in the forebrain, and regulates neuroepithelial cells in the dorsal telencephalon (Fotaki, Price and Mason, 2011; Quinn *et al*., 2009). In addition to SHH, gradients of FGF8 are essential for the development of medial and lateral forebrain areas (Zelarayan *et al*., 2007), while FGF10 is particularly important for the development of certain neuronal subtypes within the hypothalamus and interacts with SHH signaling pathways regulating forebrain development (Abler, Mansour and Sun, 2009).

To induce differentiation, we cultured our NRSCs for a total of 8 days (**Fig. 4A**), beginning with exposure to SAG (Sonic Agonist) and recombinant SHH to activate the SHH signaling pathway and promote ventralization (Belgacem *et al*., 2016). FGF8 and FGF10 were included to enhance regional patterning, guiding the cells toward a diencephalic identity (Treier *et al*., 2001). On day 4, we inhibited Notch signaling by adding DAPT (tert-butyl (2S)-2-[[(2S)-2-[[2-(3,5- difluorophenyl)acetyl]amino]propanoyl]amino]-2-phenylacetate) to facilitate neurogenesis (Louvi and Artavanis-Tsakonas, 2006). Neurotrophic factors BDNF and GDNF were administered from day 8 to support further maturation (**Fig. 4A**).

**Figure 4.**
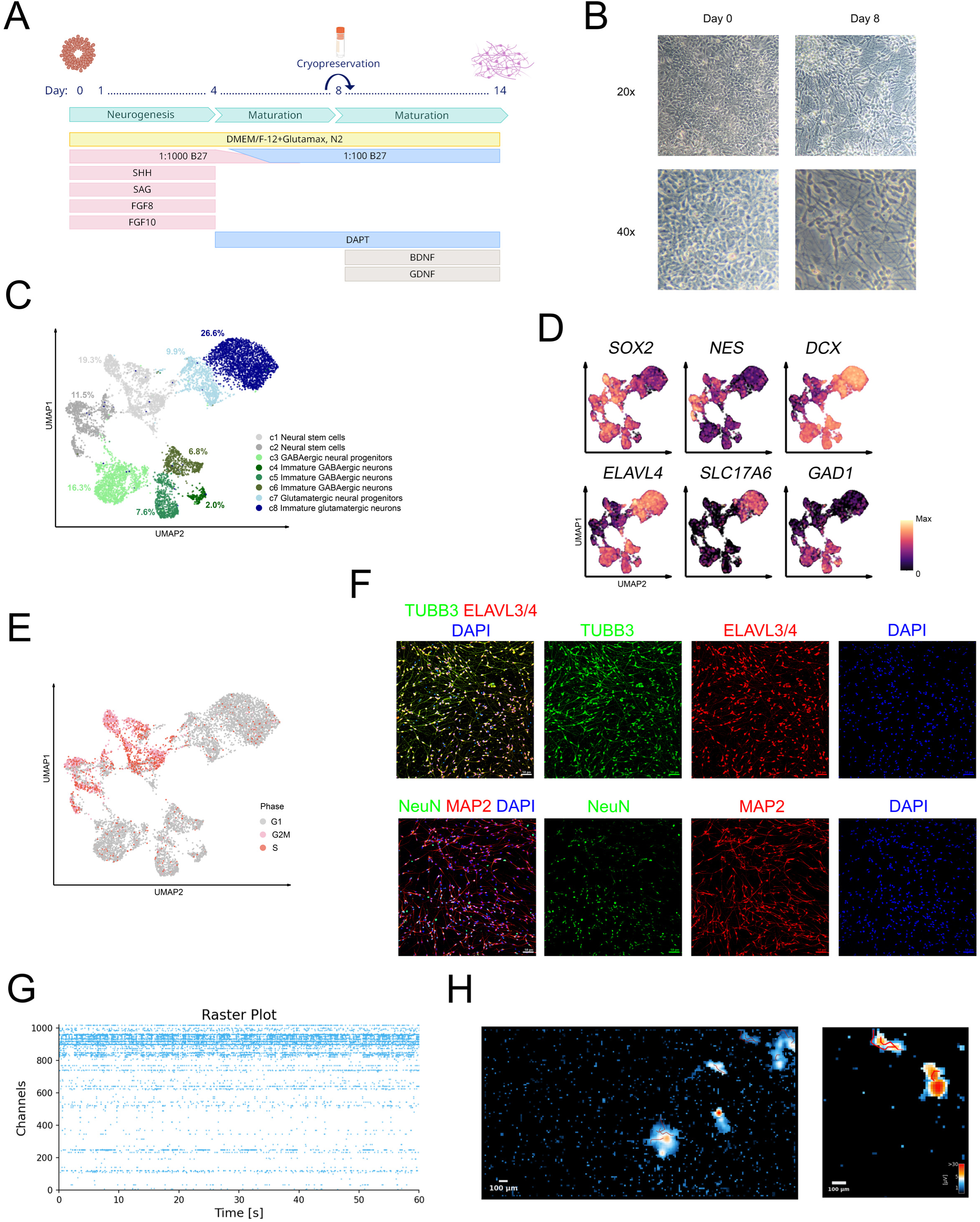
Differentiation of NRSCs into neurons. **(A)** Schematic representation of protocol for generating neurons from NRSCs. **(B)** Representative brightfield images of generated neurons captured on day 0 and 8 in 20x magnification and 40x magnification. **(C)** UMAP based on scRNA-seq of NRSC-derived neurons reveal 2 trajectories of differentiating neuronal populations. Annotation is based on known markers expressed within each cluster. scRNAseq performed after 8 days of differentiation, with day 0 corresponding to the NRSC stage. The percentage of cells in each cluster is shown color coded in the plot. **(D)** UMAP visualizing the expression levels of NSC markers; *SOX2* and *NES*, neural progenitor marker; *DCX*, pan-neuronal marker; *ELAVL4*, glutamatergic neuron marker; *SLC17A6*, and GABAergic neuron marker; *GAD1*. **(E)** UMAP visualizing the cell-cycle phase of the neurons. **(F)** Representative immunofluorescence images of NRSC-derived neurons on day 14 (with NRSC corresponding to day 0). They are positive for pan-neuronal markers TUJ1, ELAVL3/4, NeuN and MAP2. Images are taken in 20X magnification, scalebar = 50 µm. **(G)** Raster plot of spontaneous spiking by NRSC-derived neurons on HD-MEA chip at day 110 (with NRSC stage corresponding to day 0). Recording time = 60 seconds. Each dot corresponds to a spike. **(H)** Axon tracking reveal placement and axon morphology of NRSC-derived neurons on HD-MEA chip at day 110 (with NRSC stage corresponding to day 0). Higher magnification is shown to the right. Color-coding indicates recorded µV. Red lines show reconstructed axon morphology. Scalebar = 100 µm.

By day 8, significant morphological changes were observed and cells adopted common neuronal features with long axons observable via brightfield imaging (**Fig. 4B**). These immature neurons could be dissociated into single cells and cryopreserved, with an average viability above 80% upon thawing (**Suppl. Fig. 1B**).

scRNA-seq performed on day 8 characterized the cellular identities of our cryopreserved immature neurons after thawing. Analysis of 8,375 cells revealed the presence of immature glutamatergic and GABAergic neurons in our cultures (**Fig. 4C**). Predominantly, we identified proliferating neural progenitor cells (c1 and c2), marked by neural stem cell markers (NES) and proliferation markers (*MKI67* and *PCNA*) (**Suppl. Fig. 5A**). Cluster c3 represented neural progenitors positive for *STMN2*, *ELAVL4*, and *GAD1*, which transitioned into immature GABAergic neurons in clusters c4, c5, and c6. The GABAergic neurons expressed classic markers such as *DLX2*, *GAD1/2*, and *SLC32A1* (Virolainen *et al*., 2012; Zhou, Risold and Alvarez-Bolado, 2021) (**Suppl. Fig. 5B**). Furthermore, c7 and c8 indicated a trajectory toward immature glutamatergic neurons, marked by *STMN2*, *ELAVL4*, *RELN*, *ONECUT1*, and the classic glutamatergic marker *SLC17A6* (Nakamura *et al*., 2005) (**Suppl. Fig. 5C**). Notably, *RELN* has been shown to support glutamatergic synapse formation in the developing brain (Sasaki *et al*., 2014). Thus, by day 8, we observed a culture predominantly consisting of three major cell populations: *NES^+^MKI67^+^* proliferating neural progenitors and two lineages of *DCX^+^* progenitors differentiating into *ELAVL4^+^GAD1^+^* GABAergic and *ELAVL4^+^SLC17A6^+^* glutamatergic neurons (**Fig. 4D**). This was corroborated by cell cycling analysis, where proliferating neural progenitors (c1 and c2) remained in a proliferative stage, while neurons exited the cell cycle and were in the G1 phase (**Fig. 4E**).

The neuronal identity and morphology of the differentiated cells were further validated by immunofluorescence, which confirmed positive expression of pan-neuronal markers TUBB3, ELAV3/4, NeuN, and MAP2 (**Fig. 4F**). We also observed neuronal populations expressing markers associated with developing hypothalamic neurons (Kim *et al*., 2020; Zhou, Risold and Alvarez-Bolado, 2021) such as *ONECUT2*, *LHX1/2/5/9*, *KLF7*, *FOXP2*, *MEIS1*, and *SIX3* in the glutamatergic populations (clusters c7 and c8, **Suppl. Fig. 6A**), and *DLX6-AS1*, *DLX1/2*, and *ARX* in the GABAergic populations (clusters c4, c5, and c6, **Suppl. Fig. 6B**).

Overall, the scRNA-seq data provided a comprehensive characterization of the differentiated neurons, underscoring the capacity of our NRSCs to generate diverse neuronal populations.

To validate the functionality of our NRSC-derived neurons, we employed a high-density multi-electrode array system. On average, 5.1% (N=3) of electrodes were active from day 30 to day 110, with clusters exhibiting high firing rates indicative of active neurons (**Fig. 4G**). Continuous spontaneous firing of the neurons was observed over time (**Fig. 4H**). An axon tracking assay revealed four neurons on the chip, sharing a total axon length of 477.95 µm and conducting signals at a velocity of 0.504 m/s. Collectively, these results demonstrate the functional characteristics of our NRSC-derived neural progenitors.

In summary, our findings substantiate the potential to generate diverse and functional neuronal populations from our NRSCs. The successful differentiation and detailed characterization of these neurons highlights the applicability of NRSCs for *in vitro* production of functional neurons.

### NRSCs retain the capacity to differentiate into astrocytes and oligodendrocytes

NSCs are defined by their ability to self-renew and differentiate into the three primary neural lineages: neurons, astrocytes, and oligodendrocytes. The capacity for glial differentiation is a crucial criterion for validating multipotency.

To derive astrocytes from the NRSCs, we adopted a protocol based on Tcw et al. (2017), which identifies a commercial medium as an effective inducer of astrocyte differentiation from neural progenitor cells. Following this protocol, NRSCs were cultured in the commercial medium for 50 days, starting from the NRSC stage on day 0, leading to efficient conversion into astrocytes (**Fig. 5A**).

**Figure 5.**
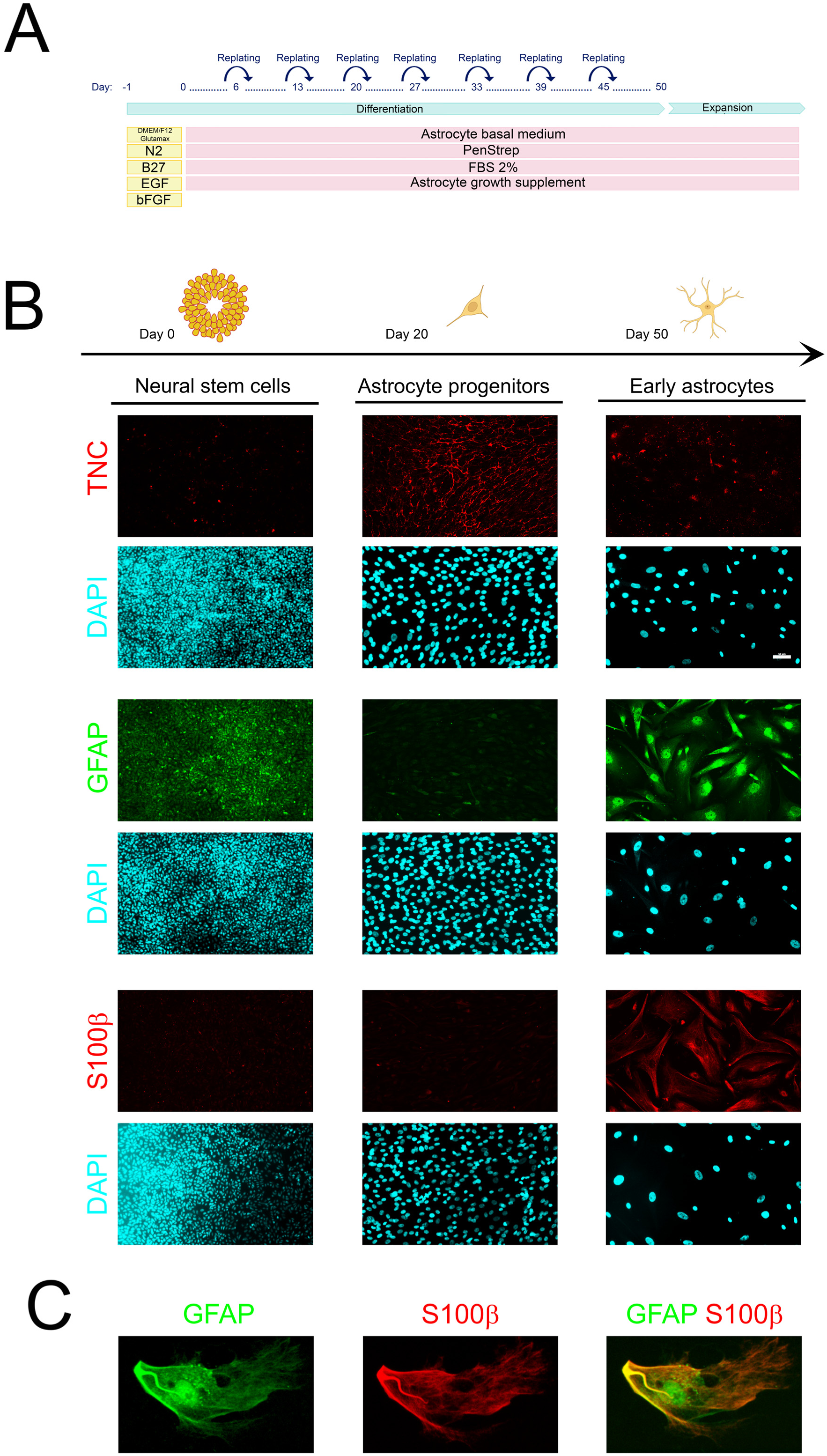
Differentiation of NRSCs into astrocytes. **(A)** Schematic representation of protocol for generating astrocytes from our NRSCs. **(B)** Representative immunofluorescence images of Astrocytes positive for TNC on day 20 and GFAP and S100β on day 50 of differentiation. Day 0 corresponds to the NRSC stage. Images are taken in 20X magnification, scalebar = 50 µm. **(C)** Higher magnified representative image of Astrocyte markers GFAP and S100β on day 55 of differentiation.

During differentiation, the astrocytes gradually acquired a star-shaped morphology with minimal branching over the 50-day period (**Suppl. Fig. 7A**). A transient expression of Tenascin-C (TNC) was noted, with TNC^+^ cells observed at day 20, which diminished substantially by day 50 (**Fig. 5B**), reflecting a common pattern in astrocyte differentiation (Faissner, Roll and Theocharidis, 2017). By day 50, we detected the astrocyte-specific markers GFAP and S100β (**Fig. 5B**), and the fibrillar morphology characteristic of more mature astrocytes (**Fig. 5C**).

Similarly, we assessed the potential for oligodendrocyte differentiation using a protocol adapted from Gorris et al. (2015) (**Fig. 6A**). This multi-stage differentiation protocol initiated with the activation of the SHH signaling pathway using SAG, promoting oligodendrocyte lineage commitment (Wang and Almazan, 2016; Santos *et al*., 2019). This was followed by the elevation of cAMP levels with Forskolin to enhance the maturation of oligodendrocyte progenitor cells (Raible and McMorris, 1989), alongside exposure to growth factors that support proliferation and survival.

**Figure 6.**
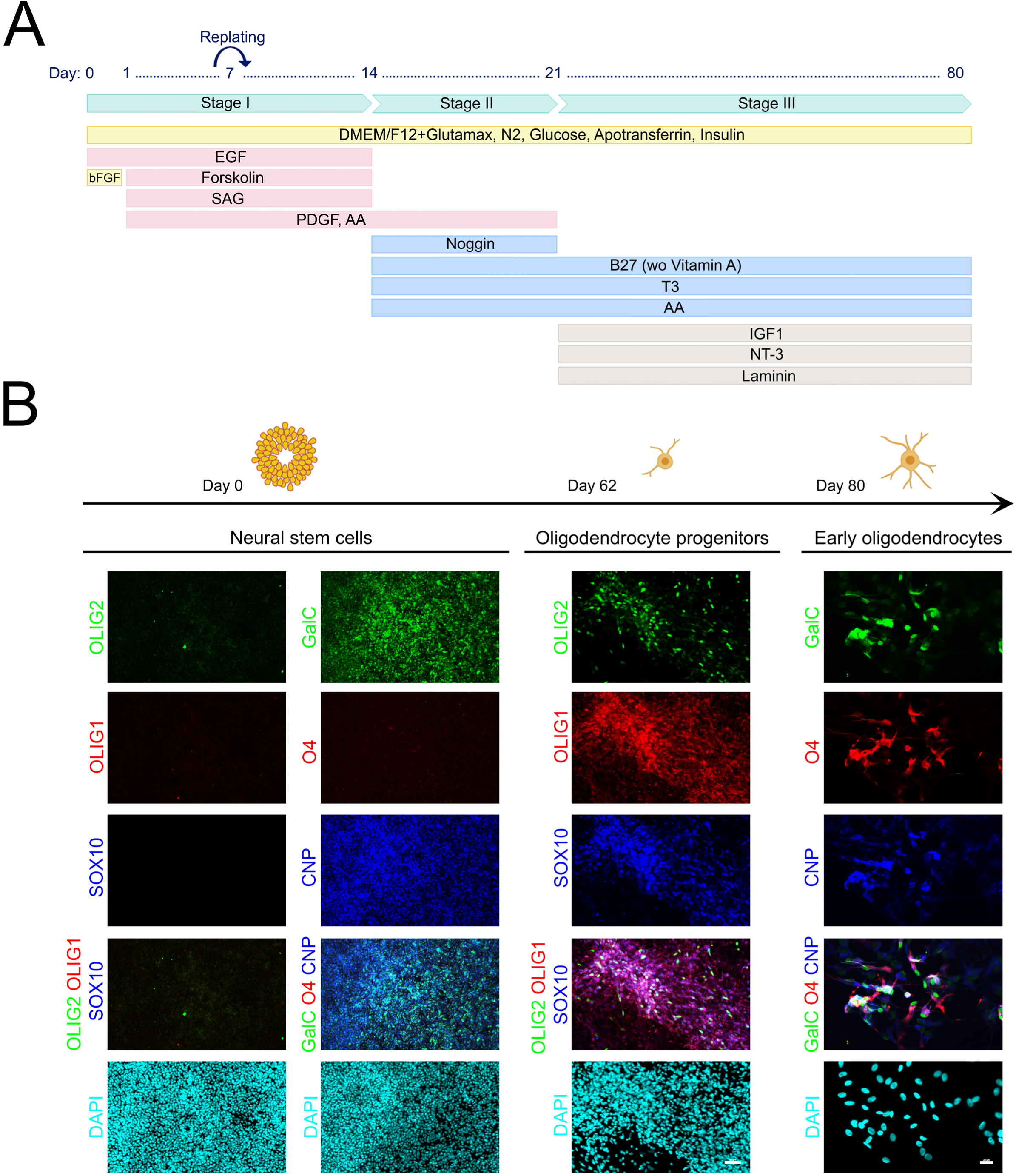
Differentiation of NRSCs into oligodendrocytes. **(A)** Schematic representation of protocol for generating oligodendrocytes from our NRSCs. **(B)** Representative immunofluorescence images of oligodendrocytes markers OLIG1, OLIG2, SOX10, O4, GALC, CNP. Images from day 0 and 62 are taken in 20X magnification, scalebar = 50 µm. Images from day 80 are taken in 40x magnification, scalebar = 20 µm.

Beginning on day 14, Noggin was introduced to inhibit the BMP pathway, which is known to facilitate oligodendrocyte differentiation. Additionally, we supplemented the culture with triiodothyronine (T3) and ascorbic acid (AA) to promote myelination and maturation (Santos *et al*., 2019). One week later, neurotrophin 3 (NT-3) and insulin-like growth factor 1 (IGF-1) were added to further stimulate myelination, along with laminin to support maturation and maintenance (Santos *et al*., 2019). The maturation of oligodendrocyte progenitor cells continued until day 80 (**Fig. 6A**).

Morphological changes were monitored throughout this differentiation process. By day 80, characteristic features of oligodendrocytes, such as small cell bodies and multiple processes, were evident (**Suppl. Fig. 7B**). The presence of oligodendrocyte-specific markers was evaluated, revealing clusters of cells positive for OLIG1/2 and SOX10 by day 62, alongside more mature markers like O4, GalC, and CNP by day 80, confirming the generation of oligodendrocytes in our cultures (**Fig. 6B**).

In summary, these results demonstrate the multipotency of our NRSCs and their ability to differentiate into glial cells, underscoring their potential applicability in disease modeling and drug screening.

## Discussion

In this study, we present a new protocol for deriving homogeneous populations of expandable NRSCs. The NRSCs generated through our approach retain multipotency and stability over multiple passages. We confirm their multipotency by differentiating them into functional neurons, astrocytes, and oligodendrocytes, showcasing their potential for diverse applications and potential for *in vitro* disease modeling, drug screening, and therapeutic interventions.

Our method addresses several limitations inherent in current NSC protocols, particularly the reliance on manual selection of neural rosettes, which represent challenges for scaling and manufacturing (Zhang *et al*., 2001; Elkabetz *et al*., 2008; Koch *et al*., 2009; Bohaciakova *et al*., 2019). Recent developments aimed at eliminating manual selection, have often resulted in heterogeneous populations or instability during passaging (Ebert *et al*., 2013; Wen and Jin, 2014; Fedorova *et al*., 2019). However, by integrating single SMAD inhibition with dynamic suspension culture, our protocol effectively removes the need for manual selection while maintaining high purity and stability, thereby streamlining the process for multiple applications. This approach facilitates a future transition into bioreactors to scale up the process.

Previous studies have indicated that NSCs tend to lose their regional identity *in vitro* over time (Elkabetz *et al*., 2008; Koch *et al*., 2009; Ebert *et al*., 2013). Notably, Koch et al. (2009) demonstrated a consistent decline in the expression of anterior markers such as OTX2 and FOXG1 after 2-10 passages. In contrast, our monitoring of FOXG1, OTX2, and PAX6, which are markers associated with anterior brain regions, showed stable expression for at least 12 passages, suggesting that our NRSCs maintain regional identity over time (**Fig. 2B**).

The observed decrease in PAX6 mRNA expression, approximately 25% from passage 2 to passage 8, juxtaposed with stable protein levels detected via flow cytometry and immunofluorescence, may suggest a complex regulatory mechanism driving *PAX6* expression. One possible explanation is the phenomenon of negative autoregulation of PAX6, where increased levels of PAX6 protein feedback inhibit its own mRNA expression (Manuel *et al*., 2007). This autoregulatory feedback loop can lead to a scenario where mRNA levels diminish while protein levels remain elevated due to translation efficiency or stability (Hsieh and Yang, 2009). As reported previously, during neurogenesis, PAX6 protein levels may not correlate linearly with mRNA levels, (Hsieh and Yang, 2009; Li *et al*., 2023).

Characterization of our NRSCs revealed high expression levels of genes associated with stemness and proliferation. However, we also encountered minor lineages of neural progenitor cells, including GABAergic and glutamatergic progenitors, as well as glial progenitors. While the NRSCs exhibited significant stemness, the differentiating cells were enriched in genes related to differentiation and maturation.

NSCs are inherently localized along the neural tube of the developing brain, each displaying distinct regional identities crucial for brain patterning and the formation of specific neuronal and glial cell types. Studies have indicated that rosette-forming NSCs, when cultured *in vitro*, can exhibit a re-specification from anterior identities toward more caudal fates, simulating the natural progression of neural tube morphogenesis (Sasai and De Robertis, 1997; Elkabetz *et al*., 2008; Koch *et al*., 2009; Ebert *et al*., 2013; Meinhardt *et al*., 2014). For example, Elkabetz et al. describe how NSCs derived from embryonic stem cells undergo differentiation into a variety of region-specific neuronal fates, but they remain partially limited in their differentiation potential without appropriate instructive cues (Elkabetz *et al*., 2008). This process highlights the intrinsic variability among NSCs, which can potentially lead to instability in the derived neurons and glial cells, presenting significant challenges for *in vitro* and therapeutic applications.

In contrast to this inherent variability, our protocol has yielded a population of highly pure FOXG1^+^OTX2^+^TLE4^+^SOX5^+^ forebrain NRSCs that exhibit remarkable regional stability across multiple passages. This retained stability throughout cell passaging represents a novel characteristic of our NRSCs, potentially offering a more consistent platform for differentiation into specific neuronal lineages (Li *et al*., 2006; Lu, 2012). However, our observations need further research to elucidate the mechanisms underlying this stability, as our current insights are limited to 12 passages, and to better understand the transcriptional and environmental factors that may contribute to these observations.

The multipotency of NRSCs is a promising asset, as demonstrated by our ability to differentiate these cells into both neuronal and glial lineages, essential for developing cell-based models and therapies. Following the findings of Julia et al. (2017), we exposed NRSCs to a commercial medium for 50 days, generating GFAP+ and S100β+ astrocytes. Moreover, we adapted an established multi-stage protocol from Gorris et al. (2015) to differentiate NRSCs into lineage-specific oligodendrocytes marked by O4, GalC, and CNP. Additionally, our NRSCs differentiated into early glutamatergic and GABAergic neurons, validated by scRNA-seq and functionality assessments using multi-electrode arrays. Our neuronal differentiation protocol aimed to generate forebrain neural progenitors. Our results suggest that we generated hypothalamic neural progenitors, denoted by markers such as *DLX6-AS1*, *ARX*, *GAD2*, *FOXP2*, *LHX2*, and *SCL17A6*, though our current findings are restricted to early neural progenitors. Future studies should focus on extended differentiation and comprehensive late-stage characterization to further explore hypothalamic neuronal specifications. Given the limited existing protocols for hypothalamic neuronal differentiation (Merkle *et al*., 2015; Ozaki *et al*., 2022), our NRSCs might provide a stable progenitor stage for generating hypothalamic neurons with reduced batch-to-batch variability. Future research should prioritize refining our protocol and further investigating the differentiation potential of NRSCs into specific neuronal subtypes.

Our NRSCs exhibit considerable differentiation potential, successfully generating pure populations of TUBB3+ neurons and early glial progenitor cells, including GFAP+ astrocytes and O4+ oligodendrocytes. This ability is critical, considering the potential of NSCs in regenerative medicine and their capacity to contribute to repair processes post-injury, underscoring their utility in therapeutic applications for neurodegenerative diseases and other neurological disorders (Jimenez- Vergara *et al*., 2023).

In summary, our results present an optimized protocol for generating highly pure NRSC populations, eliminating the need for manual rosette selection. Over multiple passages, the NRSCs consistently exhibited positivity for NSC markers such as NES and SOX2, with minimal presence of contaminating cells. Furthermore, they retained their differentiation capability into astrocytes, oligodendrocytes, and functional neurons. The protocol presented here for generating NRSC populations provides a reliable framework for advancing NSC applications in disease modeling and drug screening, as well as facilitating the exploration of their therapeutic potential in neurological diseases. This work addresses a critical gap in current methodologies, promoting the large-scale generation of pure NSC populations essential for future research and therapeutic interventions.

## Acknowledgements

We thank Frederik Møller Nielsen for support with cell culture experiments. We would like to express our gratitude to Carla Azevedo, Xiaogang Guo, and Ricardo Vieira for their invaluable discussions and support throughout this work.

## Authors contribution

SEDF and JCV conceived the study and designed the experiments. KHL and EJ contributed with the generation of scRNAseq data and analyses. CB contributed with the initial development of the differentiation protocol. SMJ contributed with flow cytometry expertise. AW contributed with the imaging platform analyses and cell culture support. SEDF performed most of the experiments. SEDF and JCV wrote the manuscript. MK, NSC and JCV supervised the project. All authors edited and approved the paper.

## Materials and methods

### Human Embryonic Stem Cell culture

The studies outlined in this manuscript were conducted in accordance with ethical guidelines and are covered by the following Danish ethical permits: H-20056277 and H-23039171 (De Videnskabsetiske Komiteer for Region Hovedstaden) for forebrain neural differentiation. hESCs lines (NOVOe001-A and E1C3 NN GMP0050E1C3) were used for several experiments conducted in this study. hESCs were cultured in Nutristem® (Sartorius: 05-100-1A) supplemented with 0,2% penicillin streptomycin (Gibco: 15140-122) in plates or flasks coated with laminin-521 (Biolamina: LN521-05). Medium was changed daily, and cells were passaged every 3-4 day before reaching 90% confluency. When passaging, the cells would first be washed with PBS-/- (GibcoTM: 14190-094) and then dissociated with TrypLETM Select (GibcoTM: 12563011) for 5-7 min at 37° C. After 5-7 min, TrypLE (GibcoTM: 12563011) was removed, and cells were resuspended immediately in Nutri supplemented with 0,2% PenStrep and Y-27632 (FujiFilm: 257-00614). The cells were counted in Nucleocounter (chemometec: NC-200) and diluted appropriately and seeded at densities of 150.000-350.000 cells/cm2. The media was changed the next day to remove Y-27632 (FujiFilm: 257- 00614). Cells were kept in a humidified incubator at 37°C and 5% CO2. Daily assessments of cell morphology were conducted in brightfield microscope. Representative images of NOVOe001-A can be found in **Supplemental Figure 8**.

### Differentiation of hESCs into NRSCs

NRSCs were differentiated from hESCs by dissociating hESCs with TrypLE^TM^ select (Gibco^TM^: 12563011) and resuspending them in NutriStem (Sartorius: 05-100-1A) supplemented with Y-27632 (FujiFilm: 257-00614) and 50 µM RepSox (Tocris: 3742). In a 12-well format, 200 000 cells were seeded in each well and kept static for 24 hours. The next day the media was changed into DMEM/F12 GlutaMAX supplement (Gibco: 31331-028), supplemented with 0,2% penicillin streptomycin (Gibco: 15140-122), 1% N2 supplement CTS (Gibco^TM^: A1370701) and 50 µM RepSox (Tocris: 3742) and the cells were placed on a Multi-bio 3D mini shaker (Biosan) at 40 rpm; 15 sec. (orbital), 5 degrees; 3 sec. (vibration) and reciprocal setting, until Day 6 to form clusters. Half the media was changed daily. Clusters were collected on Day 6 and analyzed in a BioRep (BioRep diabetes: ICC4-115/230) to determine concentration and size. Then seeded in 2D on laminin-coated (Sigma: L2020) plates, where media was changed daily. At Day 10 neural rosettes would be visible. To establish a NRSC-line we passaged and cultured the cells 3-4 times in media consisting of DMEM/F12 GlutaMAX supplement (Gibco: 31331-028), supplemented with 0,2% penicillin streptomycin (Gibco: 15140-122), 1% N2 supplement CTS (Gibco^TM^: A1370701), 1‰ B27 Supplement (Gibco^TM^: A33535-01), 10 µg/L EGF (R&D systems: 236-EG) and 10 µg/L bFGF (PeproTech: 100-18B). The NRSCs could be further cultured in the same media and passaged every 3-4 days.

### Differentiation of NRSCs into Astrocytes

NRSCs were differentiated into astrocytes by seeding dissociated single cells at a density of 15 000 cells/cm^2^ on mouse laminin-coated plates in astrocyte medium (ScienCell: 1801, astrocyte medium [1801-b], 2% fetal bovine serum [0010], astrocyte growth supplement [1852] and 0,2% penicillin/streptomycin solution (0503]). When the cells reached ∼95% confluency (approximately every 6-7 days), they were split to the initial seeding density (15 000 cells/cm^2^). After 30 days of initial differentiation, astrocytes were expanded in astrocyte medium containing 2% FBS with half the media exchanged every 2 days and passaged 1:3 every week.

### Differentiation of NRSCs into Oligodendrocytes

To generate oligodendrocytes from NRSCs, a multi-stage differentiation paradigm published by Gorris et al., 2015 was adapted. In short, cells were plated at a density of 89 000 cells/cm^2^ onto mouse-laminin (Sigma: L2020) coated cell culture-dishes (1:50 dilution in PBS+/+). The day after, medium was changed to oligodendrocyte differentiation medium stage I, consisting of DMEM/F12 GlutaMAX supplement (Gibco: 31331-028), supplemented with 0,2% penicillin streptomycin (Gibco: 15140-122), 1% N2 supplement CTS (Gibco^TM^: A1370701), 120 mg/ml D-(+)-Glucose (Sigma: G8270), 200 µg/ml Apo-transferrin (Sigma: T1147), 20 µg/ml insulin (Sigma: 19278), 10 ng/ml EGF (R&D systems: 236-EG), 10 µM Forskolin (Tocris: 1099), 1 µM SAG (Sigma: 566661) and 10 ng/ml PDGF-AA (Bio-Techne: 221-AA) . On day 7 of oligodendrocyte differentiation, cultures were replated onto mouse-laminin cell culture-dishes (1:50 dilution in PBS+/+) at a density of 9000 cells/cm^2^. On day 14 of differentiation, cultures were switched to oligodendrocyte differentiation medium stage II consisting of DMEM/F12 GlutaMAX supplement (Gibco: 31331-028), supplemented with 0,2% penicillin streptomycin (Gibco: 15140-122), 1% N2 supplement CTS (Gibco^TM^: A1370701), 120 mg/ml D-(+)-Glucose (Sigma: G8270), 200 µg/ml Apo-transferrin (Sigma: T1147), 20 µg/ml insulin (Sigma: 19278), 10 ng/ml PDGF-AA (Bio-Techne: 221-AA), 200 ng/ml Noggin (R&D systems: 6057-GMP), 2% B27 Supplement (Gibco^TM^: A33535-01), 60 ng/ml T3 (Sigma: T2877) and 200 µM Ascorbic Acid (Sigma: A4403). One week later (i.e., day 21 of differentiation), medium was changed to oligodendrocyte differentiation medium stage III consisting of DMEM/F12 GlutaMAX supplement (Gibco: 31331-028), supplemented with 0,2% penicillin streptomycin (Gibco: 15140-122), 1% N2 supplement CTS (Gibco^TM^: A1370701), 120 mg/ml D-(+)-Glucose (Sigma: G8270), 200 µg/ml Apo-transferrin (Sigma: T1147), 20 µg/ml insulin (Sigma: 19278), 2% B27 Supplement (Gibco^TM^: A33535- 01), 60 ng/ml T3 (Sigma: T2877), 200 µm Ascorbic Acid (Sigma: A4403), 10 ng/ml IGF-1 (Peprotech: 100-11), 10 ng/ml NT-3 (Peprotech: 450-03) and 1 µg/ml Laminin (Sigma: L2020). Maturation continued until day 80. In all stages half media is changed every 2-3 days.

### Differentiation of NRSC into Neurons

Neurons were differentiated by dissociating and seeding 200 000 neural stem cells/cm^2^ on mouse laminin coated plates (Sigma: L2020) in DMEM/F12 GlutaMAX supplement (Gibco: 31331-028), supplemented with 0,2% penicillin streptomycin (Gibco: 15140-122), 1% N2 supplement CTS (Gibco^TM^: A1370701), 1‰ B27 Supplement (Gibco^TM^: A33535-01), 400 nM SAG (Millipore: 566661), 500 ng/ml rhSHH (R&D systems: 1845-SH), 100 ng/ml FGF8b (Miltenyi: 130-095-738), 100 ng/ml FGF10 (Miltenyi: 130-127-855) and Y-27632 (FujiFilm: 257-00614). A full media change was performed on day 1, where Y-27621 was removed. Hereafter, we added half a volume of media every day and removed all media every second day.

On day 4 the media composition was changed to DMEM/F12 GlutaMAX supplement (Gibco: 31331- 028), supplemented with 0,2% penicillin streptomycin (Gibco: 15140-122), 1% N2 supplement CTS (Gibco^TM^: A1370701) and 10 µM DAPT (Tocris: 2634), while B27 Supplement (Gibco^TM^: A33535-01) gradually increased between day 4-6 from 1000X to 100X. The neurons were cryopreserved at DIV8.

### Cryopreservation and thawing of NRSCs and neurons

The NRSCs were cryopreserved by first dissociating them into single cell suspension with Try-pLE^TM^ select (Gibco^TM^: 12563011) for 3 min and then adding defined trypsin inhibitor (Gibco^TM^: R007100) to neutralize the reaction. They were then washed, counted and centrifuged for 3 min at 300 g. Finally, they were resuspended in StemCellBanker (Amsbio: 11924) and freezed at a controlled rate in a CoolCell (Corning) in -80° C freezer. The next day they were transferred to a liquid nitrogen tank. The NRSCs were thawed at a controlled rate in a ThawStar (Biolife Solutions) and transferred to a wash of DMEM/F12 GlutaMAX supplement (Gibco: 31331-028), supplemented with 0,2% penicillin streptomycin (Gibco: 15140-122), 1% N2 supplement CTS (Gibco^TM^: A1370701) and Y-27632 (FujiFilm: 257-00614). Hereafter they were centrifuged at 300 g for 3 min and resuspended in DMEM/F12 GlutaMAX supplement (Gibco: 31331-028), supplemented with 0,2% penicillin streptomycin (Gibco: 15140-122), 1% N2 supplement CTS (Gibco^TM^: A1370701), 1‰ B27 Supplement (Gibco^TM^: A33535-01), 10 µg/L EGF (R&D systems: 236-EG) and 10 µg/L bFGF (PeproTech: 100-18B) and Y-27632 (FujiFilm: 257-00614). The NRSCs were then seeded on mouse laminin (Sigma: L2020)-coated plates at a density of 500.000 cells/cm^2^. A full media change was performed the day after thawing, where Y-27632 (FujiFilm: 257-00614) was removed from the media from the media composition. Hereafter the media was changed.

The NRSC-derived neurons were cryopreserved by adding TrypLE^TM^ select (Gibco^TM^: 12563011) for up to 10 min until single cell suspension was obtained and subsequently adding defined trypsin inhibitor (Gibco^TM^: R007100). They were then transferred to a wash and strained through a 40 µm cell strainer (Falcon^TM^: 352340), counted and centrifuged for 3 min at 300 g. Finally, they were resuspended in StemCellBanker (Amsbio: 11924) and freezed at a controlled rate in a CoolCell (Corning) in -80° C freezer. The next day they were transferred to a liquid nitrogen tank.

The neurons were thawed at a controlled rate in a ThawStar (Biolife Solutions) and transferred to a wash of DMEM/F12 GlutaMAX supplement (Gibco: 31331-028), supplemented with 0,2% penicillin streptomycin (Gibco: 15140-122), 1% N2 supplement CTS (Gibco^TM^: A1370701), 1% B27 Supplement (Gibco^TM^: A33535-01) and Y-27632 (FujiFilm: 257-00614). Hereafter they were centrifuged at 300 g for 3 min and resuspended in DMEM/F12 GlutaMAX supplement (Gibco: 31331-028), supplemented with 0,2% penicillin streptomycin (Gibco: 15140-122), 1% N2 supplement CTS (Gibco^TM^: A1370701), 20 ng/ml BDNF (Sigma: SRP3014), 20 ng/ml GDNF (R&D systems: 212-GD-01M), 10 µM DAPT (Tocris: 2634) and Y-27632 (FujiFilm: 257-00614). The neurons were seeded on plates coated with poly-D-lysine (Gibco: A38904-01) and mouse laminin (Sigma: L2020) at a density of 150.000 cells/cm^2^. A full media change was performed the day after thawing, where Y-27632 (FujiFilm: 257-00614) was removed from the media from the media composition. For further culturing half media changes were performed every second day. DAPT was removed from the media composition on day 7 after thawing.

### Immunofluorescence

Cells were washed once with PBS^+/+^(Gibco: 14040-091), fixed in 4% PFA (ThermoScientific: J61899) for 20 min at 4°C and subsequently washed twice with PBS^+/+^. The cells were then permeabilized with 0.5% Tween-20 (VWR chemicals: 663684B) and blocked with 5% Donkey serum (Jackson Labs: 017-000-121) in 0.3% Triton X-100 (J.T. Baker: 9036-19-5) in PBS^+/+^. Afterwards, cells were incubated with primary antibodies overnight at 4°C. The next day the cells were washed twice and incubated with secondary antibodies and DAPI for 45 minutes. The cells were observed in fluorescence microscope Olympus IX81 Inverted Fluorescence Phase and BioTek Cytation C10 Confocal imaging reader and images were captured with Olympus CellSens Dimensions and BioTek Gen5 Microplate reader and imager software V3.15. A detailed list of the antibodies used can be found below.

| Antibody | Species | Fluorophore | Vendor | Cat. No. | Dilution |
| --- | --- | --- | --- | --- | --- |
| NES | Rabbit | None | Atlas antibodies | HPA007007 | 1:500 |
| TJP1 (ZO-1) | Mouse | None | Invitrogen | 33-9100 | 1:200 |
| SOX2 | Goat | None | R&D systems | AF2018 | 1:300 |
| OTX2 | Goat | None | R&D systems | AF1979 | 1:500 |
| PAX6 | Sheep | None | R&D systems | AF8150 | 1:50 |
| **FOXG1** | Rabbit | None | Abcam | Ab196868 | 1:100 |
| **NeuN** | Rabbit | None | Abcam | AB104225 | 1:500 |
| **TUJ1** | Rabbit | None | Merck | T2200 | 1:2000 |
| **MAP2** | Chicken | None | Abcam | AB92434 | 1:2000 |
| **HuC/D** | Mouse | None | Invitrogen | A21271 | 1:100 |
| **TNC** | Goat | None | R&D Systems | AF3358 | 1:20 |
| **GFAP** | Rabbit | None | DAKO | ZO334 | 1:500 |
| **S100β** | Mouse | None | Merck | S2532-2ML | 1:1000 |
| **OLIG1** | Mouse | None | Merck | MAB5540 | 1:100 |
| **OLIG2** | Rabbit | None | Merck | AB9610 | 1:500 |
| **SOX10** | Goat | None | R&D systems | AF2864 | 1:20 |
| **O4** | Mouse | None | Merck | MAB345 | 1:100 |
| **GalC** | Rabbit | None | Sigma | G9152-2MI | 1:500 |
| **CNP** | Chicken | None | Merck | AB9342 | 1:1000 |
| **NANOG** | Goat | None | R&D systems | AF1997 | 1:200 |
| **OCT3/4** | Mouse | None | Santa Cruz | Sc-5279 | 1:500 |
| **Mouse IgG** | Donkey | Alexa Fluor 555 | Thermo Fisher | A31570 | 1:800 |
| **Mouse IgG** | Donkey | Alexa Fluor 488 | Thermo Fisher | A21202 | 1:800 |
| **Rabbit IgG** | Donkey | Alexa Fluor 488 | Thermo Fisher | A21206 | 1:800 |
| **Rabbit IgG** | Donkey | Alexa Fluor 555 | Thermo Fisher | A31572 | 1:800 |
| **Goat IgG** | Donkey | Alexa Fluor 555 | Thermo Fisher | A21432 | 1:800 |
| **Goat IgG** | Donkey | Alexa Fluor 647 | Thermo Fisher | A21447 | 1:800 |
| **Sheep IgG** | Donkey | Alexa Fluor 647 | Thermo Fisher | A21448 | 1:800 |
| **Chicken IgY** | Goat | Alexa Fluor 555 | Thermo Fisher | A32932 | 1:800 |
| **Chicken IgY** | Goat | Alexa Fluor 647 | Thermo Fisher | A32933 | 1:800 |

### Flow Cytometry

Cryopreserved neural stem cells at various passages were thawed and centrifuged before being resuspended in DMEM/F12 GlutaMAX supplement (Gibco: 31331-028), supplemented with 0,2% penicillin streptomycin (Gibco: 15140-122) and 1% N2 supplement CTS (Gibco^TM^: A1370701) to achieve a concentration of 1×10^6^ cells/mL. The cells were incubated with LIVE/DEAD™ Fixable Near- IR viability dye (1:1000, Invitrogen: L10119) for 15 min at room temperature, protected from light, to identify dead cells. Subsequently, the cells underwent a single wash in Wash buffer (PBS-/- (Gibco^TM^: 14190-094) supplemented with 1% bovine serum albumin (Miltenyi Biotec: 130-091-376) followed by centrifugation (400xg for 10 min). The cells were fixed and permeabilized using the Transcription Factor Buffer Set (BD Pharmingen: 562574) according to the manufacturer’s instructions. For each intracellular staining, 250 000 fixed cells were pelleted by centrifugation (800xg for 3 minutes) and then incubated with a cocktail of conjugated antibodies (see detailed list below) diluted in Perm/Wash buffer for 30 minutes at +4°C. After incubation, the cells were washed twice with Perm/Wash buffer, pelleted by centrifugation (800xg for 3 minutes), and finally resuspended in Wash buffer. Prior to acquisition on a BD FACSymphony A5 flow cytometer, the cells were filtered through a 40-µm MultiScreen-MESH Filter Plate (Millipore: MANMN4010). 20,000 live single cells were acquired per sample. Compensation was achieved using UltraComp eBeads™ Plus Compensation Beads (Invitrogen: 01-3333-42) and the ArC Amine Reactive Compensation Bead Kit (Invitrogen: A10346). The FCS files were analyzed using FlowJo software (v.10.10.0), and the population gates were set based on appropriate unstained and fluorescence-minus-one (FMO) controls.

| Antibody | Conjugation | Vendor | Cat. No. | Dilution | Cell type stained |
| --- | --- | --- | --- | --- | --- |
| **OTX2** | VioB515 | Miltenyi Biotec | 130-121-193 | 1:320 | NRSCs |
| **SOX2** | BV421 | BioLegend | 656114 | 1:130 | NRSCs/hESC |
| **PAX6** | APC | Miltenyi Biotec | 130-123-267 | 1:160 | NRSCs |
| **SOX1** | PE | Miltenyi Biotec | 130-111-043 | 1:320 | NRSCs |
| **NANOG** | AF647 | BioLegend | 674010 | 1:1000 | hESC |
| **OCT3/4** | PE | BD Biosciences | 560186 | 1:20 | hESC |
| **TRA-1-60** | BV421 | BD Biosciences | 562711 | 1:20 | hESC |
| **SSEA4** | V450 | BD Biosciences | 561156 | 1:20 | hESC |

### Single cell RNA sequencing

scRNA-seq of NSC and neuronal cultures was conducted using the 10X Genomics Chromium Single Cell 3′ Reagent Kit version 3.1 protocol. Libraries were prepared with unique molecular index (UMI) barcoding, amplified, and sequenced on an Illumina NovaSeq 6000 platform. The resulting sequence reads underwent processing using the Cell Ranger pipeline (v7.1, 10X Genomics). The preprocessing steps included quality control, alignment, filtering, barcode counting, and UMI counting. Specifically, reads were aligned to the human reference genome GRCh38 (Ensembl release 107) following Cell Ranger’s standard workflow. The resulting feature-barcode matrices were subjected to further analysis using R (v4.2) and the Seurat package (v5). Initial quality control involved filtering cells based on two criteria: (1) the number of detected genes (nFeature_RNA) within the range of 1000 to 10 000 and (2) a mitochondrial read percentage of less than 10%. For samples P2 and P8, integration was carried out to correct for batch effects utilizing Seurat’s canonical correlation analysis (CCA) approach, while NRSC samples were processed using a standard merge workflow without batch correction. Following normalization, cell cycle scoring was implemented using Seurat’s built-in cell cycle markers, with subsequent regression of cell cycle effects during data scaling to mitigate cell cycle-associated variation. Dimensionality reduction was achieved through principal component analysis (PCA), followed by UMAP for visualization. Graph-based clustering was performed to discern distinct cell populations. Differential expression analysis between clusters was conducted using Seurat’s FindAllMarkers function. Cluster identification was achieved through the integration of known literature markers and differentially expressed genes. Visualizations were created using the Seurat, scCustomize, and ggplot2 packages in R.

### High density multi-electrode array

MaxOne chips (MaxWell biosystems: PSM) were sterilized by 1% Terg-A-Zyme treatment, followed by submersion into ethanol and washing with deionized water. The chips were hereafter filled with complete culture medium and pre-conditioned for 3 days in an incubator. Before plating the cells, chips were coated with poly-D-lysine (Gibco: A38904-01) and mouse laminin (Sigma: L2020). 150000 neurons were seeded by placing 50 ul cell suspension onto the chip, followed by 60 min incubation.

60 min later, 600 µl were added of DMEM/F12 GlutaMAX supplement (Gibco: 31331-028), supplemented with 0,2% penicillin streptomycin, 1% N2 supplement CTS (Gibco^TM^: A1370701), 2% B27 (Gibco: A33535-01), 20 ng/ml BDNF (Sigma: SRP3014) and 20 ng/ml GDNF (R&D systems: 212- GD-01M). The neurons were maintained for 3 months by changing half the media every 2-3 days. Recording of electrical activity were conducted 24 hours after media change, on day 110 (with the NRSC-stage corresponding to Day 0), on a MaxOne Single-well HD-MEA system (MaxWell). Analysis was performed with the MaxLive software (MaxWell) and included activity analysis, network analysis and axon tracking analysis.

### Statistical analyses

All data were collected from at least three independent experiments and presented as mean ± standard deviation unless otherwise specified. Due to the qualitative nature of the data, no formal statistical tests were performed. The mean and standard deviation were calculated for some quantitative measurements obtained, providing a descriptive summary of the data.

### Use of generative AI and AI-assisted technologies

Generative AI tools were used by the authors to improve readability and language of the original text. No generative AI tools were used to create any of the data or to create or alter images.

---

**Supplementary figure 1.**
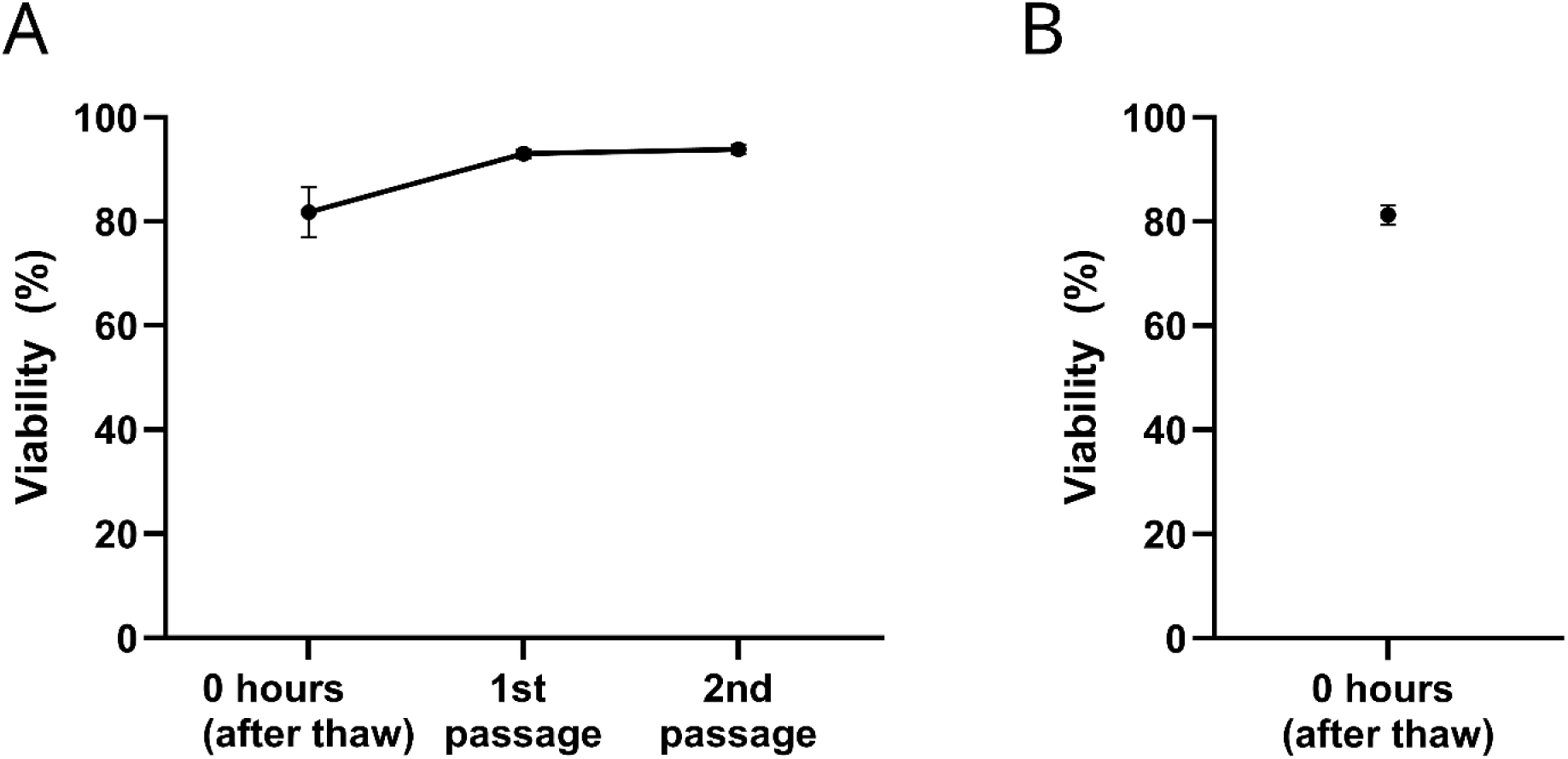
NRSCs and neurons maintain high viability after cryopreservation. **(A**) Viability of NRSCs after thawing and the 1^st^ and 2^nd^ passage after thawing. N=3. **(B)** Viability of NRSC-derived neurons after thawing. Shown as mean +/- SD, N=5.

**Supplementary figure 2.**
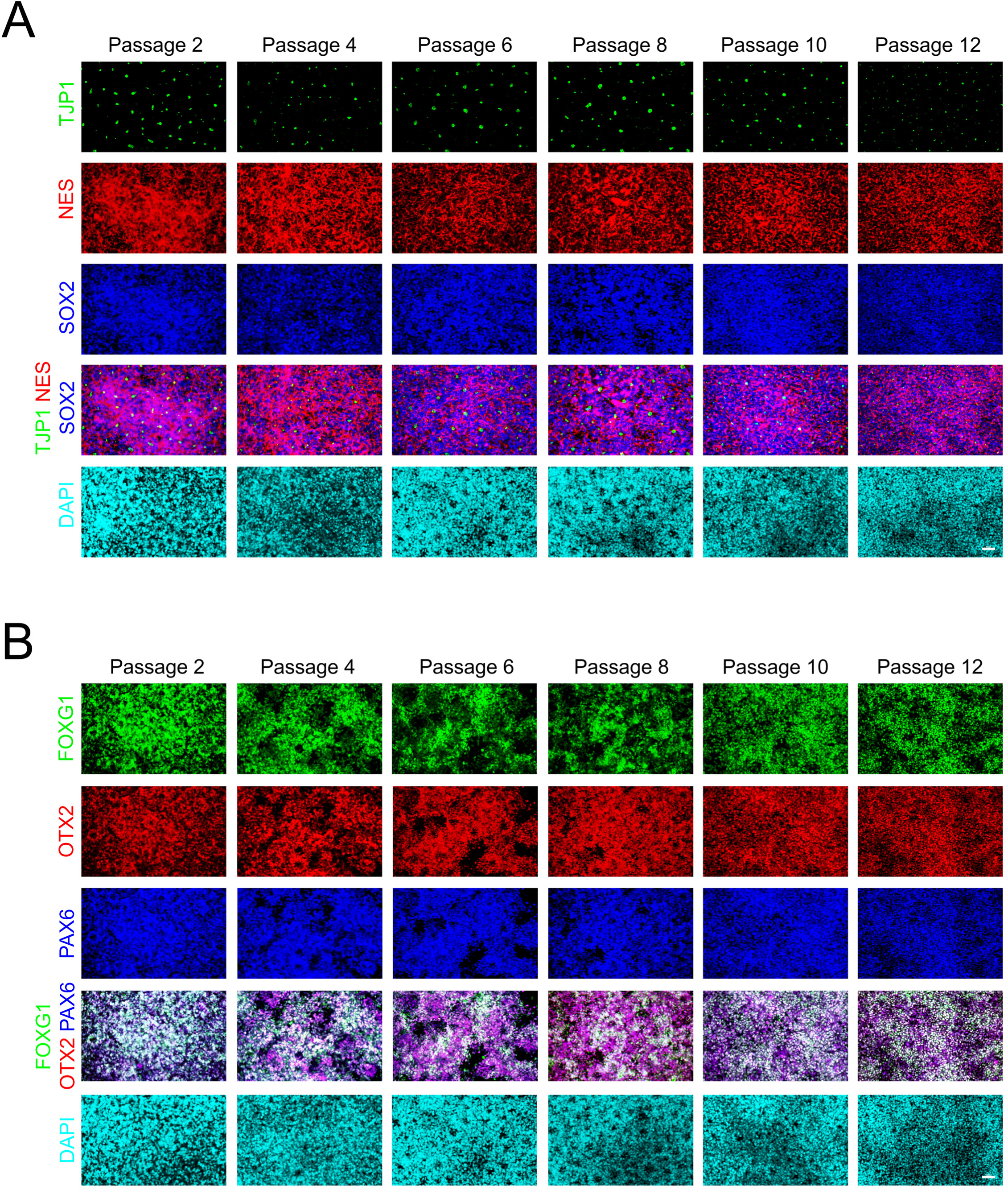
Separate NRSC-line maintain stable marker expression through extended passaging. **(A)** Representative immunofluorescence images show cells positive for NSC markers NES and SOX2 in at passage 2, 4, 6, 8, 10 and 12. TJP1 is used to highlight the lumen of neural rosettes. All images are taken in 20X magnification, scalebar = 50 µm. The different passages are imaged at different timepoints. **(B)** Representative immunofluorescence images show NRSC positive for regional forebrain markers FOXG1 and OTX2. And NSC marker PAX6. All images are taken in 20X magnification, scalebar = 50 µm. The different passages are imaged at different timepoints

**Supplementary figure 3.**
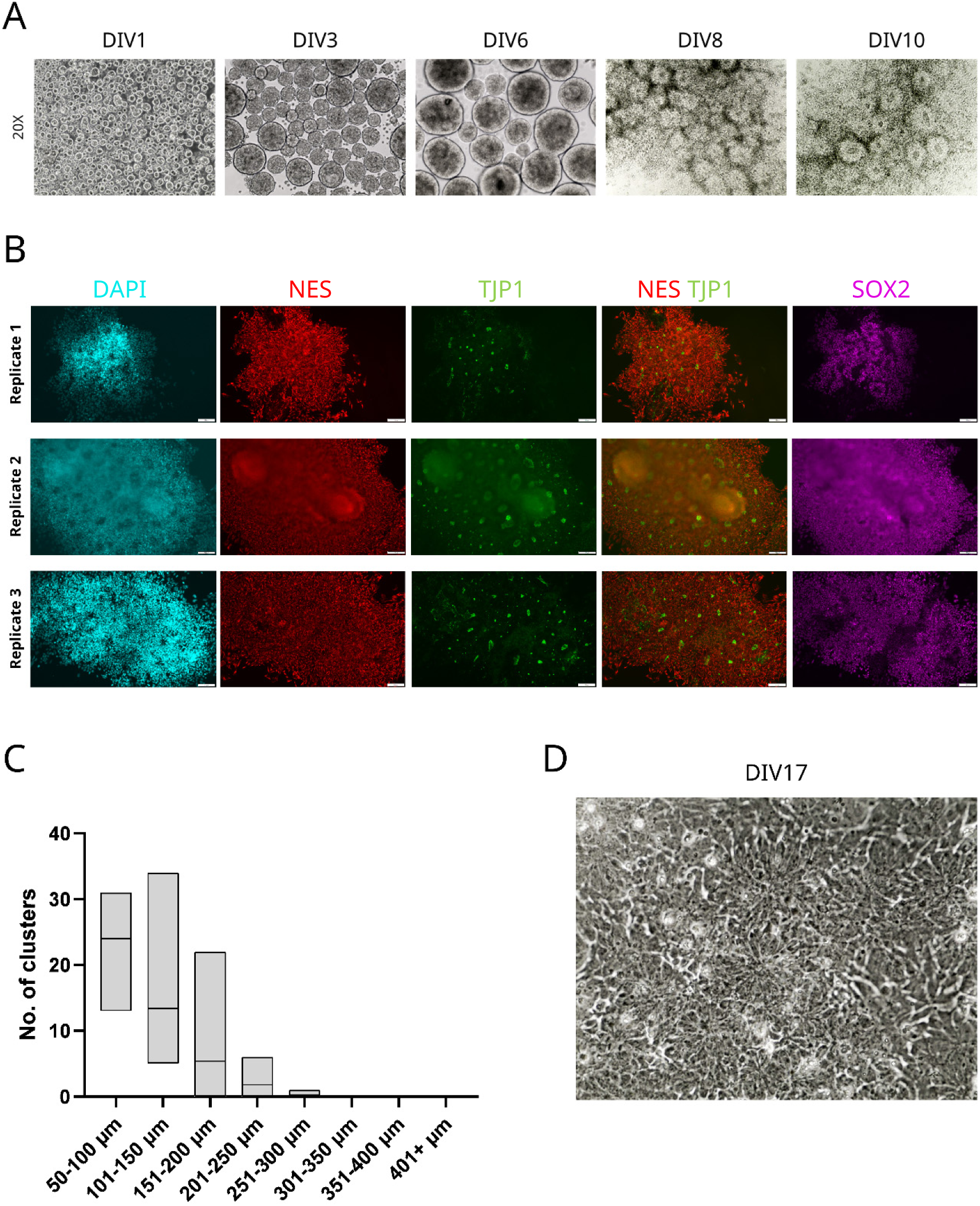
Generation of NRSCs from alternative cell line. **(A)** Representative brightfield images captured on DIV 1, 3, 6, 8 and 10 in 20x magnification. **(B)** Representative immunofluorescence images of DIV 10 NRSCs positive for common NSC markers NES and SOX2. The formation of neural rosettes is visualized by the redistribution of TJP1 into the lumen of the rosettes. Scalebar = 100 µm. **(C)** Size distribution of differentiating clusters measured on DIV 6 before seeding in 2D. Box represents minimum and maximum values, line is mean, N=5. **(D)** Representative brightfield image captured on DIV 17 in 40x magnification, revealing the formation of rosettes.

**Supplementary figure 4.**
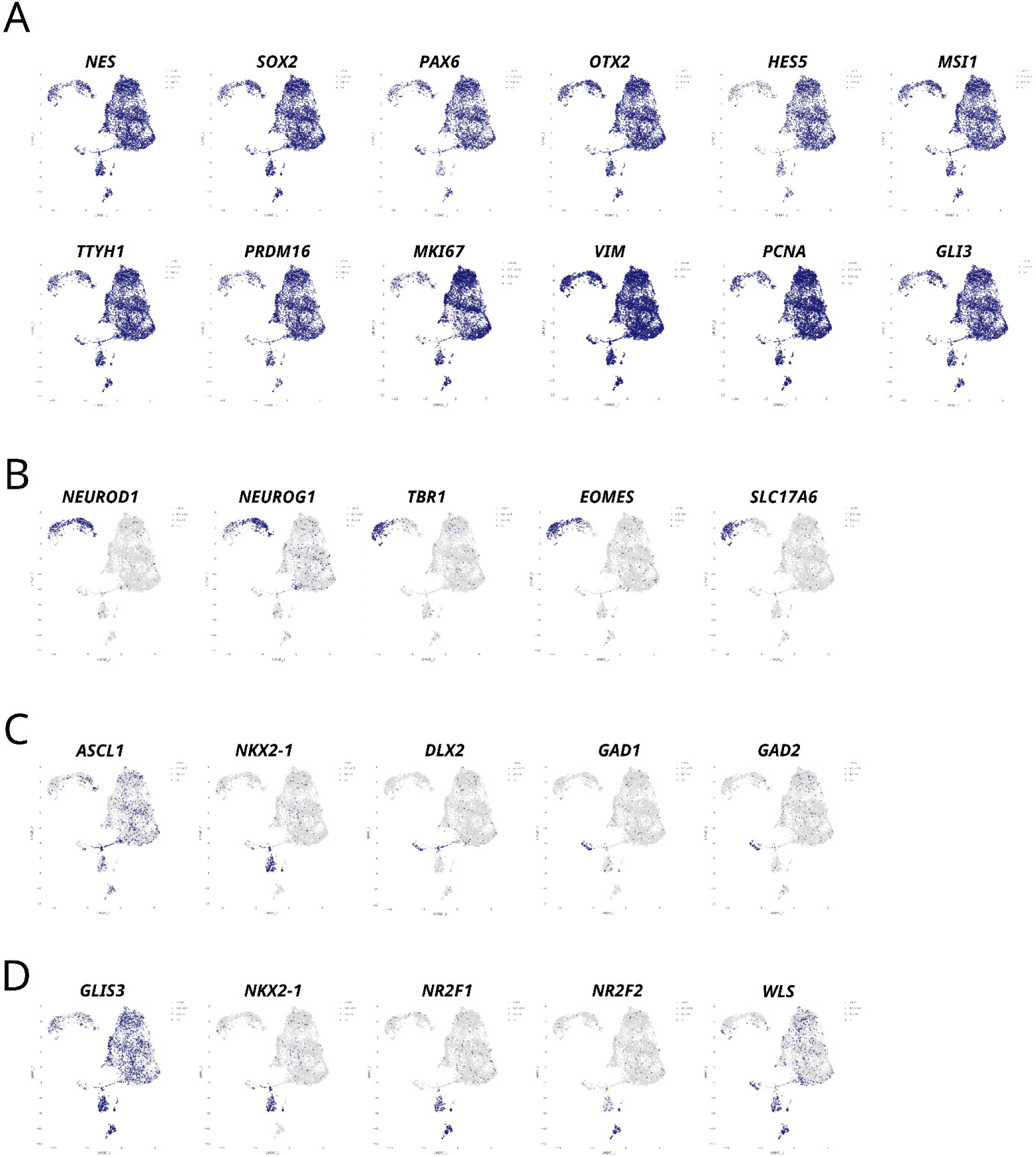
scRNA-seq analyses indicate high cell purity of NRSCs. **(A)** UMAP visualization based on scRNA-seq of NRSCs show the expression levels of NSC markers; *NES, SOX2, PAX6, OTX2, HES5, MSI1, TTYH1, PRDM16, MKI67, VIM*, *PCNA* and *GLI3* **(B)** UMAP visualization based on scRNA-seq of NRSCs show the expression levels of glutamatergic neural progenitor markers; *NEUROD1, NEUROG1, TBR1, EOMES* and *SLC17A6*. **(C)** UMAP visualization based on scRNA-seq of NRSCs show the expression levels of GABAergic neural progenitor markers; *ASCL1, NKX2-1, DLX2, GAD1* and *GAD2*. **(D)** UMAP visualization based on scRNA-seq of NRSCs show the expression levels of glial progenitor markers; *GLIS3, NKX2-1, NR2F1, NR2F2* and *WLS*.

**Supplementary figure 5.**
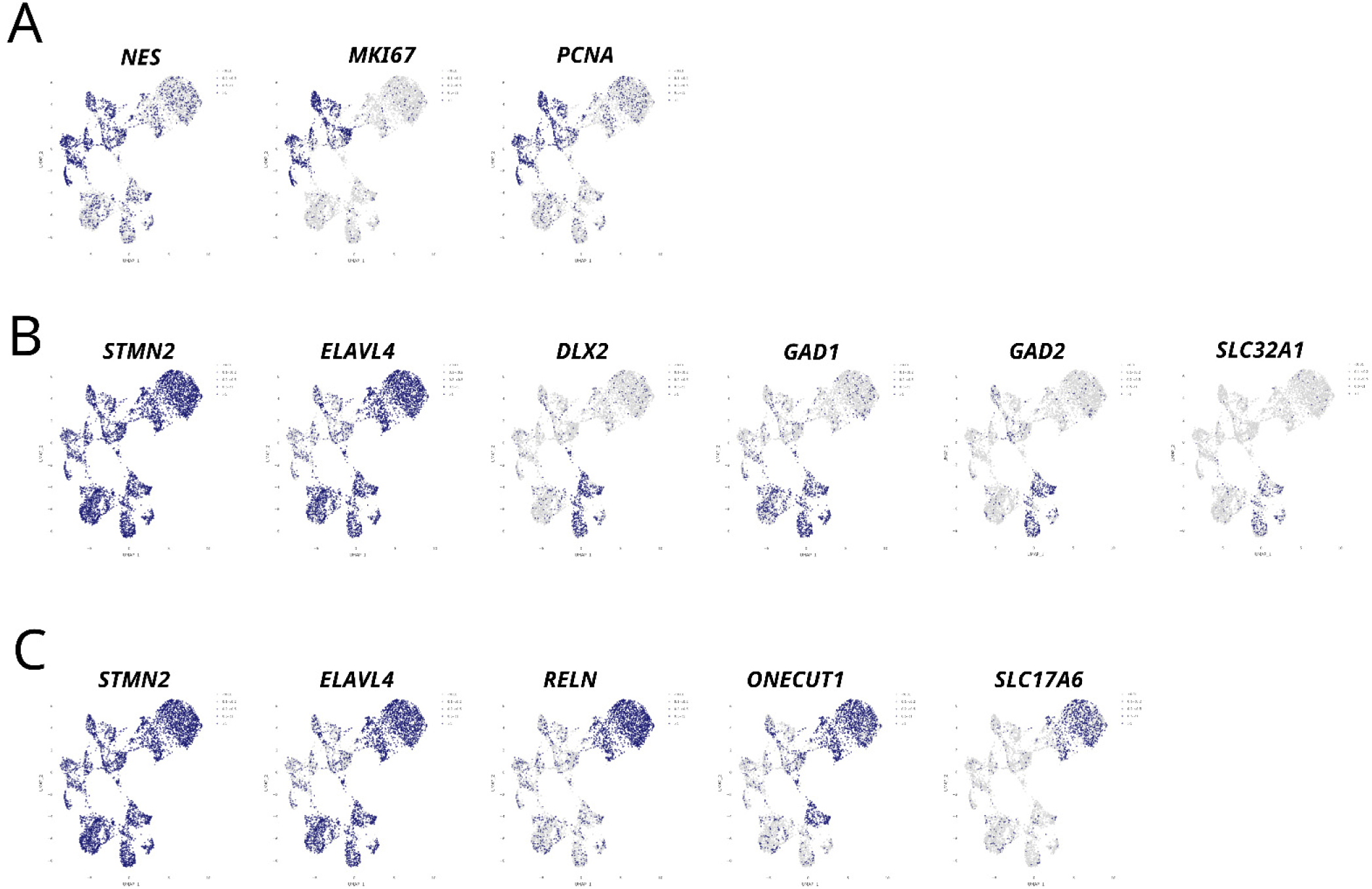
scRNA-seq analyses reveal neuronal differentiation capacity of NRSCs. **(A)** UMAP visualization based on scRNA-seq of NRSC-derived neurons show the expression levels of NSC marker; *NES* and proliferation markers: *MKI67* and *PCNA*. **(B)** UMAP visualization based on scRNA-seq of NRSC-derived neurons show the expression levels of pan-neuronal markers; *STMN2* and *ELAVL4* and GABAergic neural progenitor markers; *DLX2, GAD1, GAD2* and *SLC32A1*. **(C)** UMAP visualization based on scRNA-seq of NRSC-derived neurons show the expression levels of pan-neuronal markers; *STMN2* and *ELAVL4* and glutamatergic neural progenitor markers; *RELN, ONECUT1* and *SLC17A6*.

**Supplementary figure 6.**
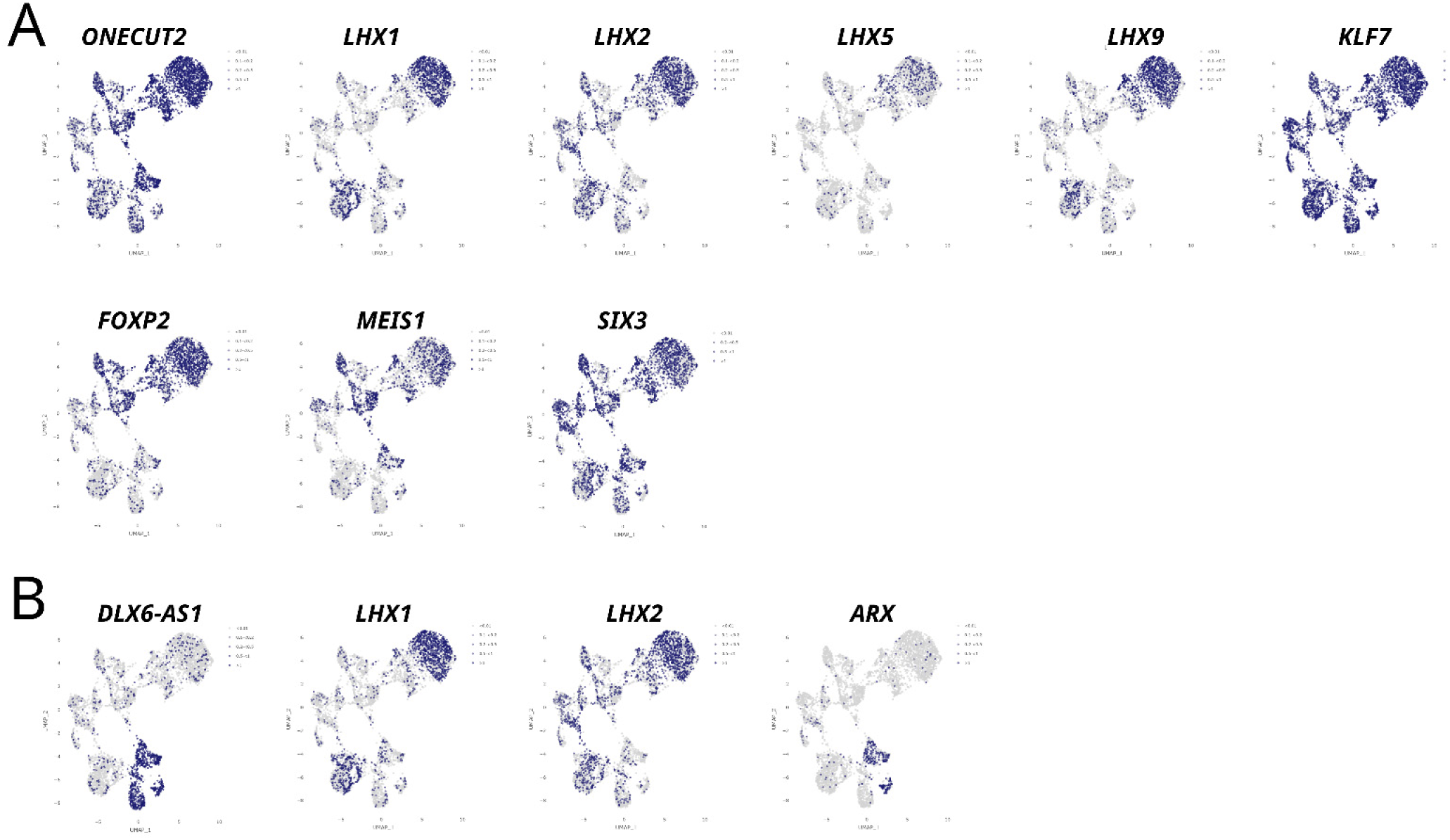
Early hypothalamic identity of NRSC-derived neurons. **(A**) UMAP visualization based on scRNA-seq of NRSC-derived neurons show the expression levels of glutamatergic early hypothalamic markers; *ONECUT2, LHX1/2/5/9, KLF7, FOXP2, MEIS1 and SIX3*. **(B)** UMAP visualization based on scRNA-seq of NRSC-derived neurons show the expression levels of GABAergic early hypothalamic markers; *DLX6-AS1, LHX1/2* and *ARX*.

**Supplementary figure 7.**
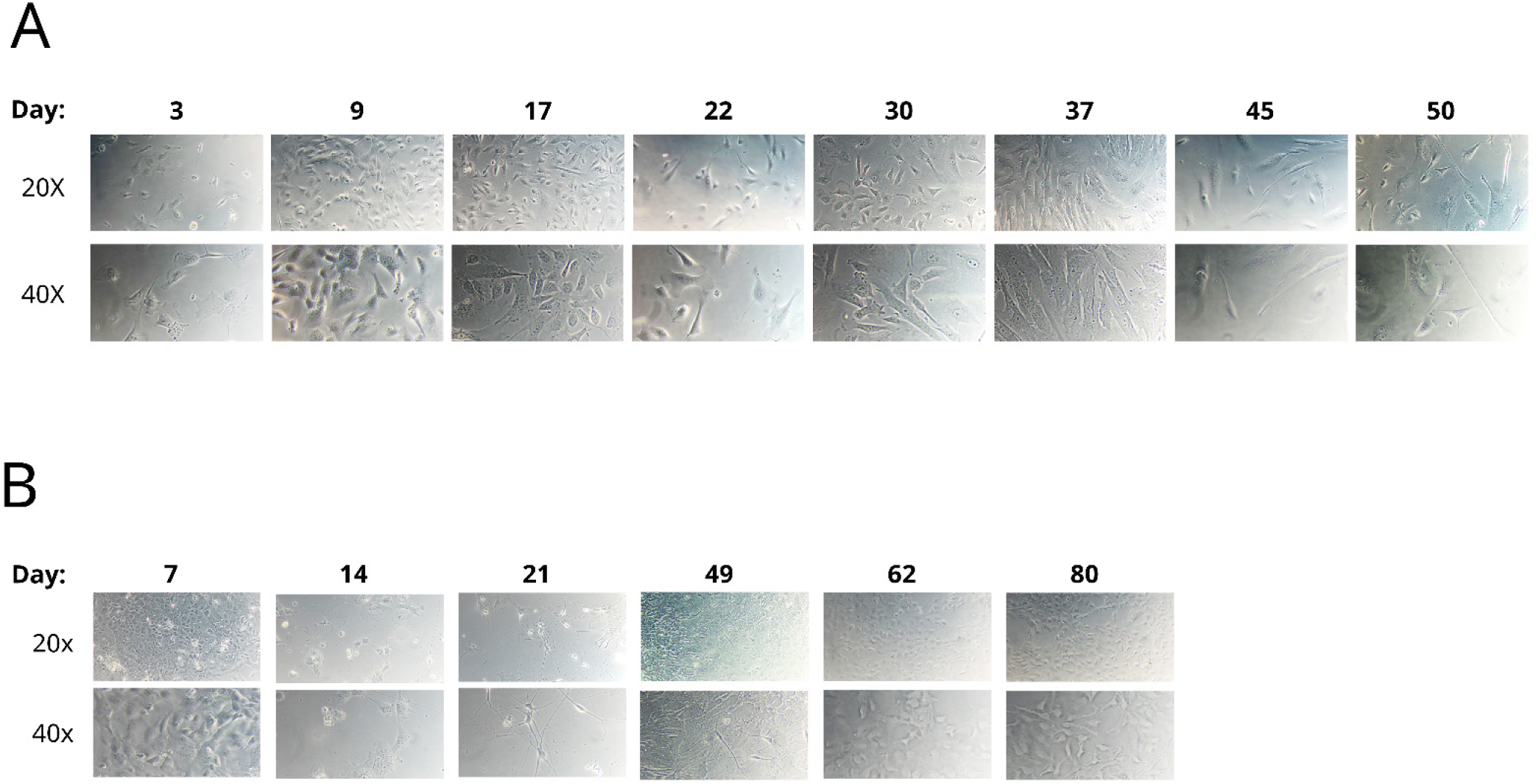
Morphological development during glial differentiations. **(A)** Representative brightfield images of astrocyte differentiation captured on DIV 3, 9, 17, 22, 30, 37, 47 and 50 in 20x magnification and 40x magnification. DIV 0 corresponds to NRSC-stage. (B) Representative brightfield images of oligodendrocyte differentiation captured on DIV 7, 14, 21, 49, 62, and 80 in 20x magnification and 40x magnification. DIV 0 corresponds to NRSC-stage

**Supplementary figure 8.**
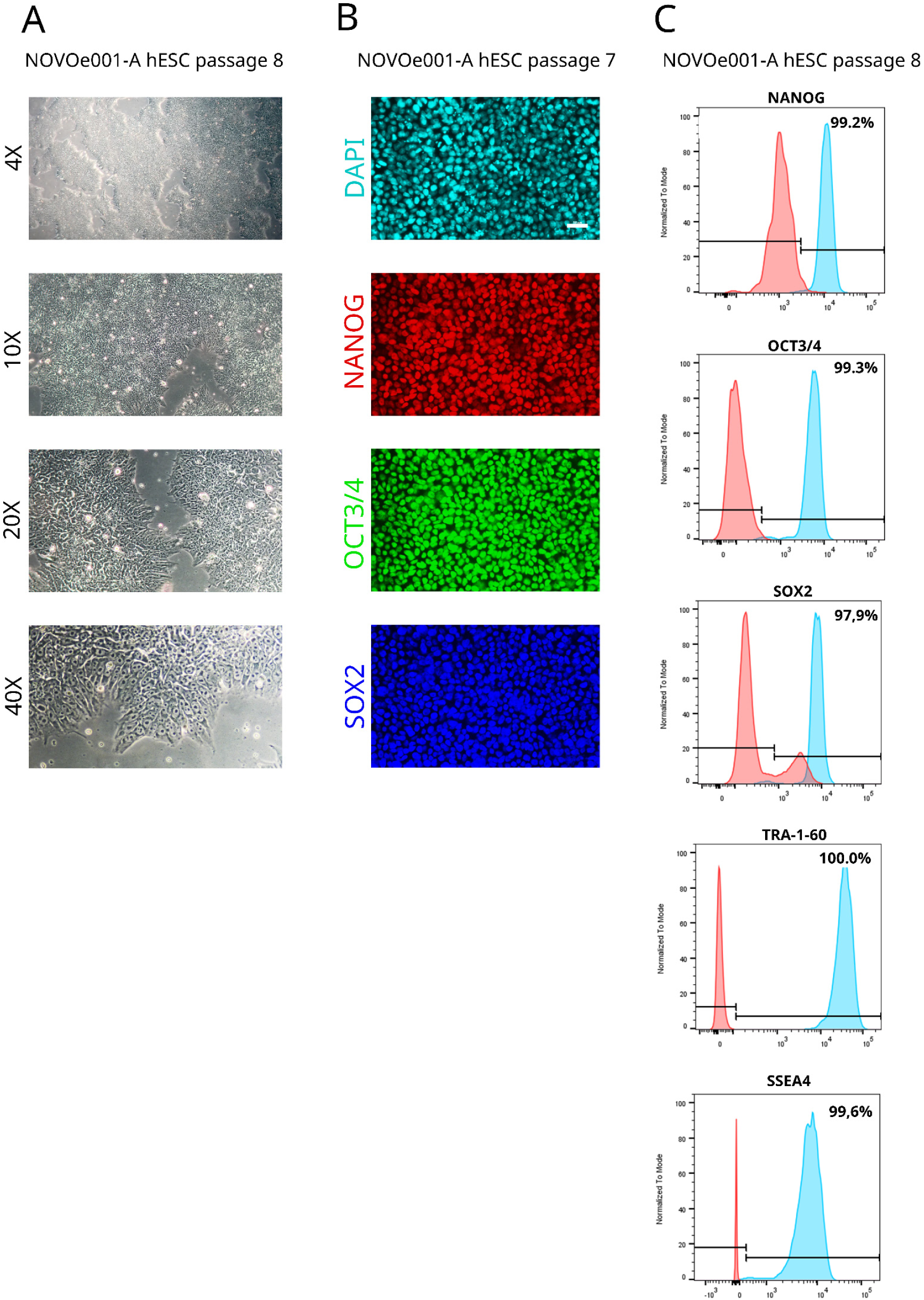
hESC line characterization. **(A)** Representative brightfield images of NOVOe001-A hESC line at passage 8 showing clear hESC morphology with large cell-bodies, dark nucleus and spiky edges. Taken in 4X, 10X, 20X and 40X magnification. **(B)** Representative immunofluorescence images of NOVOe001-A hESC line at passage 7 showing positive for pluripotency markers NANOG, OCT3/4 and SOX2. Scalebar = 50 µm. **(C)** Flow cytometry histrograms at passage 8 showing positive expression for pluripotency markers NANOG, OCT3/4, SOX2 and TRA-1-60.

